# TL1A overexpression in Crohn’s Disease and mice alters Paneth cells and microbiota promoting ileal inflammation

**DOI:** 10.1101/2025.03.02.641061

**Authors:** Yuefang Ye, Shyam K. More, Hussein Hamade, Erica E. Alexeev, Yosuke Shimodaira, Anna Y. Blackwood, Dalton T. Stamps, Jordan H. Miller, Jay P. Abraham, Lisa S. Thomas, Sofi L. Castanon, Hannah Q. Estrada, Kotaro Kumagai, Alka A. Potdar, Talin Haritunians, Emebet Mengesha, Kolja Wawrowsky, Shrikant Bhute, Richard S. Blumberg, Dermot P. B. McGovern, R. Balfour Sartor, David Q. Shih, Robert J. Barrett, Noam Jacob, Jonathan P. Jacobs, Stephan R. Targan, Kathrin S. Michelsen

## Abstract

Paneth cells regulate host-microbial homeostasis and defects in autophagy and host defense pathways have been associated with inflammatory bowel diseases (IBD). Genetic variants in TL1A (*TNFSF15*) and its receptor DR3 (*TNFRSF25*) have been associated with IBD. TL1A expression is increased in IBD patients, particularly in TL1A risk allele carriers. However, effects of TL1A on Paneth cells, resident microbiota, and development of ileitis remain unknown. TL1A overexpression in mice induces Paneth cell hyperplasia and morphological abnormalities preceding the development of ileitis. In Crohn’s disease (CD) patients, ileal TL1A expression was associated with abnormal Paneth cell phenotypes. We confirmed direct effects of TL1A on Paneth cells in human iPSC-derived human intestinal organoids and mouse Paneth cell-enriched organoids. Resident microbiota was required for TL1A-mediated Paneth cell dysfunction, and ileitis. *Tl1a-tg* mice were enriched in short chain fatty acid-producing bacteria and the metabolite acetate. Acetate supplementation in WT or *Tl1a-tg* mice caused ileal inflammation, suggesting that acetate is sufficient to cause ileitis. DR3-deficiency in Paneth cells resulted in Paneth cell abnormalities and microbiome composition changes. Our findings provide a mechanistic link between overexpression of TL1A in CD patients, Paneth cell dysfunction, and enrichment of acetate-producing bacteria and acetate that promotes ileal inflammation.

**Brief Summary:** Overexpression of TL1A drives Paneth cell dysfunction in Crohn’s Disease and mice leading to microbial and metabolomic changes that promote small bowel inflammation.

## Introduction

Inflammatory bowel diseases (IBD) are heterogeneous chronic inflammatory disorders caused by dysregulated immune responses to intestinal microbiota in genetically predisposed individuals (1). The two major types of IBD are Crohn disease (CD), which can affect any segment of the gastrointestinal tract with distinctive pathological features such as patchy transmural inflammation, relative sparing of the rectum and intestinal fibrostenosis, and ulcerative colitis (UC), which is limited to the colonic mucosa. Genome-wide association studies have identified multiple loci involved in immune responses, host-microbial interactions, and cellular processes such as autophagy and endoplasmic reticulum (ER) stress responses to be associated with susceptibility to IBD. Variants in the genes *TNFSF15* (encoding the protein TL1A) and its receptor *TNFRSF25* (encoding the protein DR3) are associated with susceptibility to and severity of IBD (2–4). TL1A expression is elevated in inflamed intestinal areas of CD patients, and CD patients with a *TNFSF15* risk haplotype have elevated peripheral TL1A expression and develop more severe disease (5, 6). Mouse models of constitutive TL1A overexpression that mimic IBD patients with *TNFSF15* risk polymorphisms cause spontaneous ileitis and fibrostenosing disease (6–8). Recent phase II clinical trials using neutralizing anti-TL1A antibodies have shown promising results in patients with UC (9). While most of the pro-inflammatory effects of TL1A have been attributed to its effects on innate and adaptive immune cells, the role of TL1A on non-immune cells is less defined (6–8, 10, 11). We have previously demonstrated that TL1A induces fibroblast activation and development of fibrosis in TL1A-overexpressing mice and CD patients with *TNFSF15* risk variants develop more frequently stricturing disease (6, 7, 12). However, the role of TL1A on other non-immune cells in the mucosa that may contribute to the development of IBD such as Paneth cells have not been investigated.

Paneth cells are specialized epithelial cells primarily located in the crypts of the small intestine that regulate host-microbial homeostasis by secreting antimicrobial peptides. Several IBD risk polymorphisms manifest in Paneth cell defects, particularly in the host defense, autophagy, and ER stress response pathways, and have been associated with the development of intestinal inflammation in CD patients and mouse models and were shown to be the origin of intestinal inflammation (13–16). Paneth cell dysfunction leads to intestinal dysbiosis in CD patients and in mouse models (17). As a consequence of intestinal dysbiosis, CD patients also present with a shift in microbial metabolites that includes a reduction of protective short-chain fatty acids, particularly butyrate, and an increase of primary bile acids and sphingolipids (18–20). Particularly secondary bile acids found to be increased after high fat diet consumption are central in complex feedback mechanisms involving Paneth cell dysfunction that leads to microbiota dysbiosis, enrichment in secondary bile acids which exerts direct toxic effects on Paneth cells and thereby exacerbate Paneth cell dysfunction (19, 21).

Genetic risk factors in these pathways that determine susceptibility to IBD are also modulated by environmental factors. Environmental triggers that cause Paneth cell dysfunction include cigarette smoking, diet, or infection with murine norovirus (19, 22, 23). In addition to environmental factors, host derived cytokines also affect Paneth cell function. In particular, cytokines associated with mucosal immune responses to pathogens such as IL-22, and IL-17 induce anti-microbial peptides in Paneth cells directly or via promoting the commitment of intestinal stem cells to the secretory lineage (24–27). IFN-γ has also been shown to be an important trigger of Paneth cell degranulation and release of anti-microbial peptides (28). Identifying additional genetic, microbial, and metabolomic risk factors that might contribute to Paneth cell dysfunction could help develop novel intervention strategies for CD.

Here, we identified the interaction of TL1A, Paneth cells, and resident microbiota and their metabolite acetate as drivers of the development of ileitis. We show that mucosal TL1A expression in CD patients correlates with Paneth cell abnormalities. Overexpression of TL1A in mice (*Tl1a-tg* mice) leads to Paneth cell dysfunction that precedes ileal inflammation and is at least partially dependent on the expression of the TL1A receptor DR3 on Paneth cells. Using human iPSC-derived human intestinal organoids (HIO) enriched for Paneth cells in a microfluidic device subjected to continuous laminar flow and mouse Paneth cell-enriched organoids we demonstrate that TL1A directly alters Paneth cell phenotypes. Furthermore, *Tl1a-tg* mice are enriched in acetate-producing bacteria *Bifidobacterium* and *Allobaculum* and metabolomic analysis revealed an increase of the metabolite acetate. Acetate supplementation in germ-free (GF) or SPF housed WT or *Tl1a-tg* mice caused ileal inflammation and elevated IFN-γ secretion, suggesting that acetate is sufficient to cause ileitis. Furthermore, DR3-deficiency in Paneth cells lead to changes in the ileal microbial composition. Our findings provide a mechanistic link between overexpression of TL1A in CD patients with *TNFSF15* risk variants, Paneth cell dysfunction, enrichment of acetate-producing bacteria and acetate that promote ileal inflammation. Our data have implications for how microbiota-derived acetate may elicit previously unknown pro-inflammatory effects in the context of genetic IBD risk factors.

## Results

### Paneth cell abnormalities precede the development of small intestinal inflammation in mice overexpressing TL1A

We and others have previously demonstrated that mice over-expressing TL1A (*Tl1a-tg*) develop spontaneous ileitis over time (7, 8). Histological examinations revealed increased ileal inflammation and increased numbers of goblet and Paneth cells compared to wild-type (WT) mice. However, the molecular mechanisms that lead to Paneth cell hyperplasia in *Tl1a-tg* mice have not been explored. First, we analyzed Paneth cell numbers, morphology, and the degree of ileal inflammation in WT and *Tl1a-tg* mice at different ages (Figures 1A-C). Interestingly, we observed Paneth cell hyperplasia in *Tl1a-tg* mice as early as 4 - 6 weeks of age when there were no signs of ileitis (Figures 1A, B). In the process of aging, the numbers of Paneth cells increased in both WT and *Tl1a-tg* mice but *Tl1a-tg* mice have significantly higher numbers of Paneth cells at any time-point (Figure 1A). In mice older than 6 months of age we observed ileitis in *Tl1a-tg* mice but not in WT mice (Figure 1B). Paneth cell abnormalities in the intracellular distribution of granules containing antimicrobial proteins such as lysozymes have been implicated in a subset of CD patients and mice with genetic deletions in the autophagy and ER stress pathways (13–15, 17, 29). We observed marked changes in Paneth cell morphology in 6-month-old *Tl1a-tg* mice with a dominant diffuse lysozyme staining compared to normal granule packaging in WT mice (Figure 1C). Next, we used a previously established scoring system to identify and quantify Paneth cells with abnormal lysozyme staining in these mice at different ages (22). In 4- to 6-week-old WT mice we observed Paneth cells with different degrees of granule abnormalities that shifts to a predominant D0 phenotype characterized by normal granule size, numbers, and distribution upon aging (Figures 1D-F). In contrast, 4- to 6-week-old *Tl1a-tg* mice predominantly displayed Paneth cells with D3 abnormalities (diffuse lysozyme staining) that was also apparent in older mice (Figures 1D-F). In comparison to WT mice, *Tl1a-tg* mice displayed a significantly higher degree of Paneth cell abnormalities at any age but most pronounced in older mice and these abnormalities preceded the development of ileitis (Figures 1B, D-F). These results suggest that overexpression of TL1A drives changes in Paneth cell numbers and morphology preceding the development of ileitis. Transmission electron microscopy confirmed Paneth cell abnormalities in *Tl1a-tg* mice. We observed disordered and distended ER and mitochondria and increased vesiculation in 7-month-old *Tl1a-tg* mice while WT mice displayed well organized ER structure (Figure 1G). Moreover, in *Tl1a-tg* mice we observed more Paneth cell granules with less electron dense content surrounded by expanded electron-lucent halos formed by packaging of Muc2 mucin potentially representing a degranulation or defect in the packaging of electron dense contents into granules, a phenotype that has also been observed in Paneth cells of mice deficient in the CD susceptibility gene *IRGM1* (30, 31). These data suggest that Paneth cells in *Tl1a-tg* mice have defective intracellular protein trafficking.

**Figure 1.**
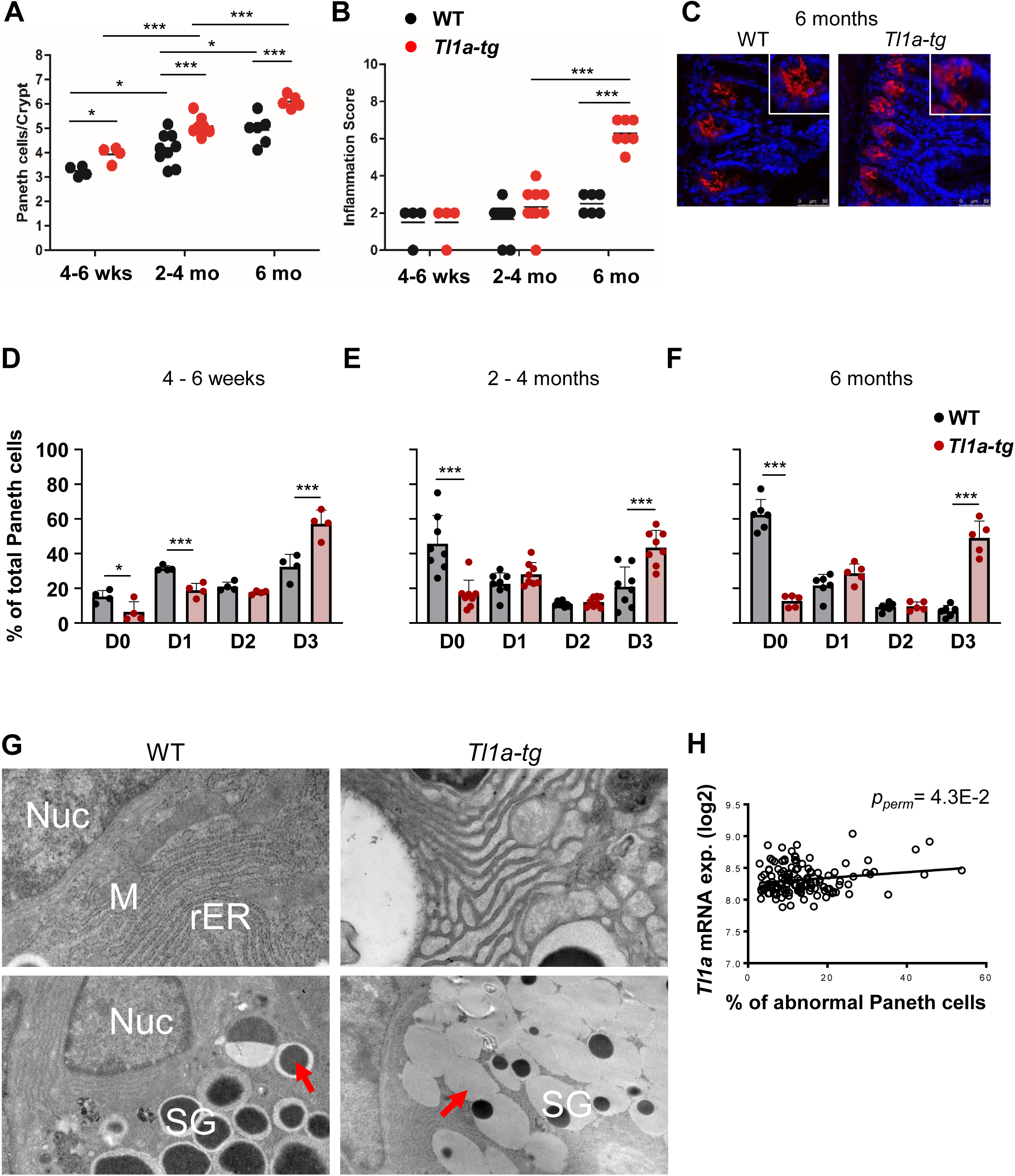
TL1A-overexpression is associated with abnormal Paneth cell morphology in mice and CD patients. **(A)** Numbers of Paneth cells in WT and *Tl1a-tg* mice at indicated time-points. **(B)** Histologic ileal inflammation score was determined by a blinded observer. **(C)** Representative lysozyme staining in 6-months-old WT and *Tl1a-tg* mice. **(D-F)** Percentage of Paneth cells with D0-3 features in *WT* and *Tl1a-tg* mice at indicated ages. (n = 4-9 mice/group). Data represent means ± SD. \**p* < 0.05; \*\**p* < 0.01; \*\*\**p* < 0.005, two-tailed Student’s *t*-test. **(G)** Representative transmission electron microscopy images of WT and *Tl1a-tg* Paneth cells. Nuc = nucleus, SG = secretory granules, M = mitochondria, rER = rough endoplasmatic reticulum. Red arrows indicate high-density secretory granules (magnification: 3,600x). **(H)** Quantitative trait associations in CD patients between small intestinal *TNFSF15* gene expression and abnormal Paneth cell phenotypes (D1-D4) using first two principal components in genotype data and gender as covariates. Associations were confirmed by permutations and reported as permuted *p*-value (p_perm) (n = 132 CD patients).

### Mucosal TL1A mRNA expression in uninflamed ilea of CD patients is associated with Paneth cell abnormalities

We previously identified a *TNFSF15* risk haplotype associated with increased expression of TL1A in peripheral monocytes (5). To determine the human relevance of our findings from murine TL1A over-expressing mice, we analyzed the degree of Paneth cell abnormalities in post-surgical tissue sections of uninflamed ilea in a cohort of genotyped CD patients (n = 132 subjects) (Figure 1H, Table 1) (32). We analyzed association of TL1A mRNA expression with Paneth cell abnormalities in this dataset of 132 CD patients. Quantitative trait associations were performed between *TNFSF15* gene expression and various Paneth cell phenotypes (D0, D1, D2, D3, D4, D1234 in %) using first two principal components in genotype data and sex as covariates. Only the associations with D1, D3 and D1234 (all abnormal Paneth cell phenotypes) were found to be significant (Figure 1H, Table 1). We confirmed the associations by permutations and report a permuted *p*-value. Addition of *ATG16L1* T300A (rs2241880) as covariate did not affect the *TNFSF15* mRNA association with Paneth cell phenotypes. In contrast, we did not observe an association of *NOD2* mRNA expression with any Paneth cell phenotypes (*p* < 0.05, data not shown). These data suggest that TL1A expression correlates with Paneth cells abnormalities in human CD and is an *ATG16L1* T300A independent association.

**Table 1:** Association between Paneth cell phenotype and mucosal TL1A expression.

| **N** | **Gene** | **PC**<br>**Phenotype** | **beta** | **p-value** | **N_perm** | **p_perm** |
| --- | --- | --- | --- | --- | --- | --- |
| 132 | TNFSF15 | D1 | 1.63 | 1.49E-03 | 10000 | **2.40E-03** |
| 132 | TNFSF15 | D3 | 4.63 | 7.14E-03 | 10000 | **9.60E-03** |
| 132 | TNFSF15 | D1234 | 6.89 | 4.35E-02 | 1000 | **4.30E-02** |

### Global gene expression analysis of Paneth cells reveals TL1A over-expression affects protein ubiquitination, cellular transports, apoptosis, and ER stress response

To determine the signaling pathways downstream of TL1A we performed RNA-Sequencing analysis on Paneth cells from WT and *Tl1a-tg* mice. In brief, we isolated Paneth cell-containing intestinal crypts by laser capture microscopy on ileal sections from 8-week-old mice at a time point when no inflammation was observed. We identified 59 genes differentially expressed in *Tl1a-tg* vs WT mice at adjusted *p* < 0.01 (Figure 2A). Enrichment and Ingenuity Pathway analysis of differentially expressed transcripts revealed significant overrepresentation of biological processes of intracellular protein traffic, protein targeting and localization (Figure 2B). These results are consistent with Paneth cell granule packaging abnormalities in *Tl1a-tg* mice (Figure 1). We also observed enrichment of transcripts encoding transporters and ligases, particularly those in protein ubiquitination pathways that are enriched in *Atg16l1^HM^*mice (22). We also observed a significant up-regulation of apoptotic genes in *Tl1a-tg* mice including Bak1 (Figures 2A, B). Next, we examined the induction of epithelial cells death in *Tl1a-tg* mice. Surprisingly, we observed a significant reduction of TUNEL^+^ cells in the crypt-base of *Tl1a-tg* mice compared to WT mice in 8-week- and 6-month-old mice (Figures 2C, D). While the number of TUNEL^+^ cells in the crypt-base increased in WT mice over time we did not observe any increase of TUNEL^+^ cells in *Tl1a-tg* mice.

**Figure 2.**
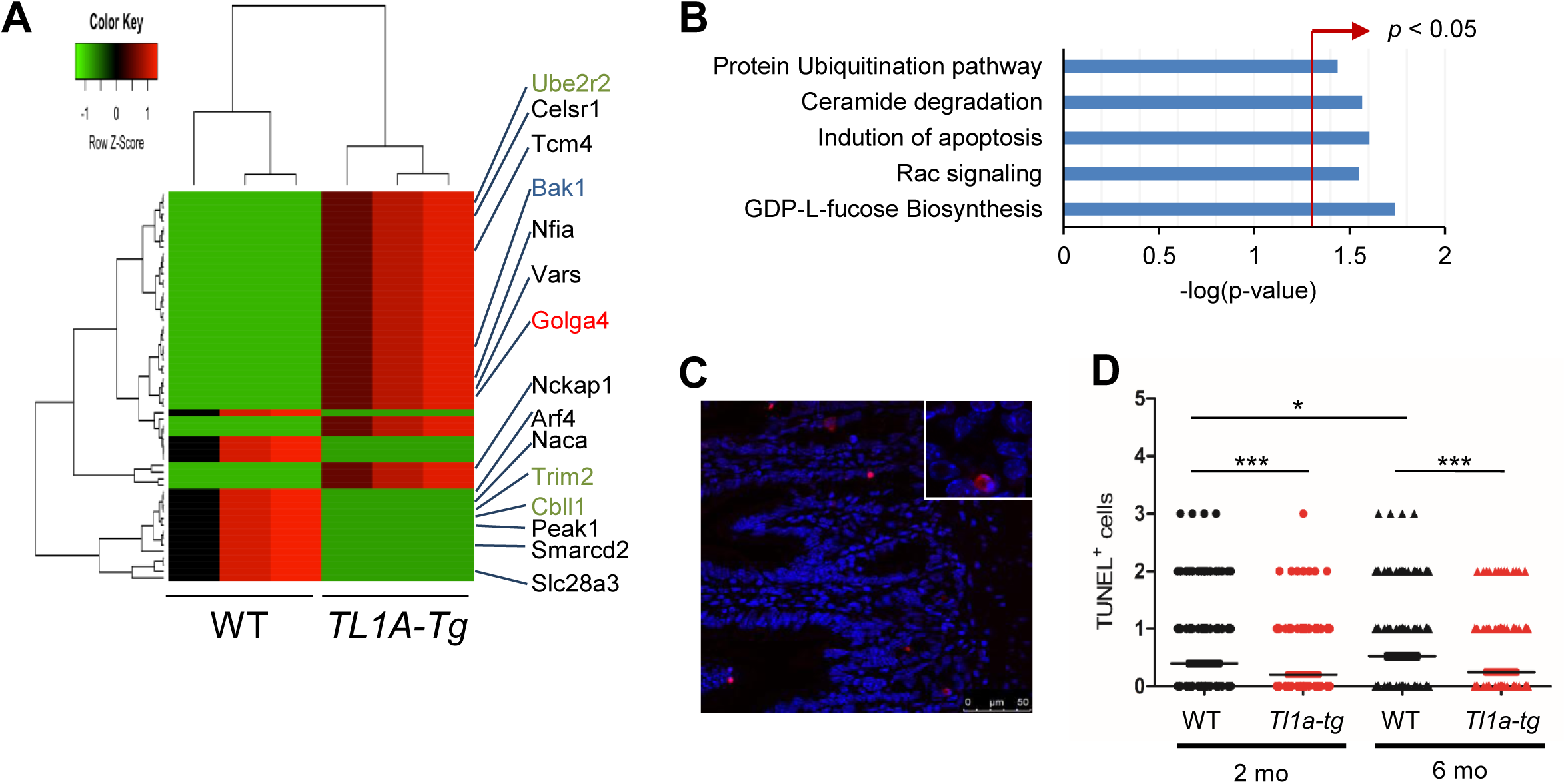
TL1A-overexpression leads to transcriptional changes in Paneth cells preceding small intestinal inflammation and is associated with abnormal Paneth cell granules, and ER stress. (**A-C**) Transcriptional profiling of small intestinal crypts of untreated 2-month-old WT and *Tl1a-tg* mice (n = 3 mice/group). (**A**) Heat map displaying RNA sequencing data of top 59 genes differentially expressed with *p* < 0.05. The dendrograms to the left and above the heat map represent hierarchical clustering of genes (rows) and samples (columns). Selected genes highlighted in red are associated with cellular transports, green with protein ubiquitination, and blue with induction of apoptosis. (**B**) Enrichment of pathways from genes differentially expressed in WT vs. *Tl1a-tg* Paneth cells. (**C**) Representative image of TUNEL staining in WT mice. (**D**) Quantification of TUNEL^+^ cells in ileal crypts of 2, and 6-month-old WT and *Tl1a-tg* mice. At least 40 crypt-villus units were quantified per mouse (n = 5-6 mice/group). Data points represent individual crypts. \**p* < 0.05; \*\*\**p* < 0.005, two-tailed Student’s *t*-test.

### Death Receptor 3 (DR3) is expressed on murine and human ileal Paneth cells

To determine if the effects of TL1A overexpression on Paneth cell hyperplasia and morphology is directly mediated, we investigated the expression of the TL1A receptor Death receptor 3 (DR3) on Paneth cells. We used single-molecule fluorescent in situ hybridization (smFISH) for DR3 and co-stained with anti-lysozyme antibodies to identify Paneth cells within ileal crypts (Supplemental Figures 1A, B). We observed expression of DR3 mRNA in lysozyme-positive Paneth cells in WT mice (Supplemental Figure 1A). We did not observe expression of DR3 in *Dr3^-/-^* mice demonstrating specificity of our DR3 probe (Supplemental Figure 1B). We confirmed expression of DR3 in ileal tissue sections from CD patients and observed distinct expression on lysozyme-positive Paneth cells and enterocytes (Figure 3A).

**Figure 3.**
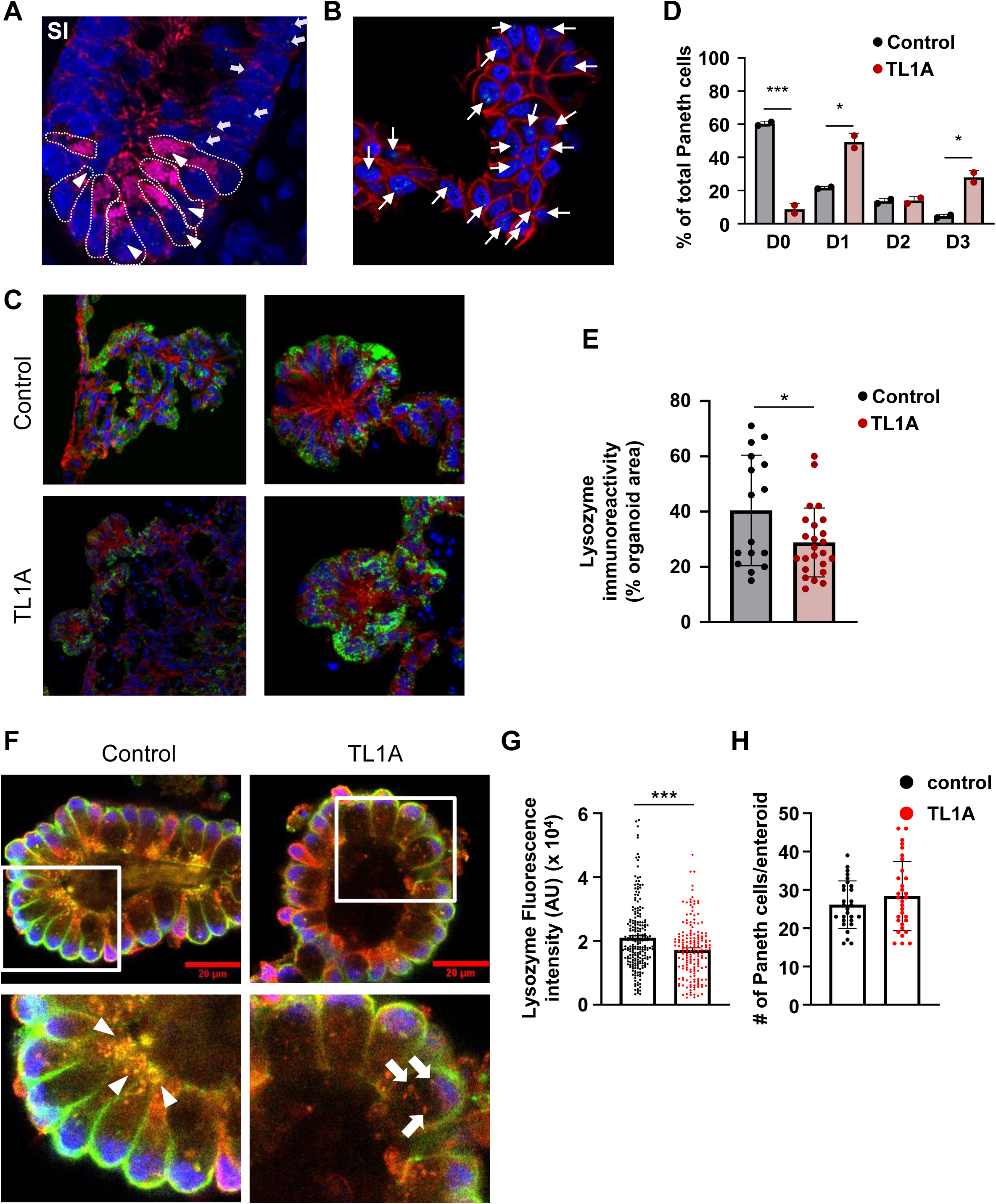
TL1A drives abnormal Paneth cell development in human and mouse intestinal organoid derived Paneth cells. (A) Expression of human DR3 in non-inflamed small intestinal tissue sections from CD patient. DR3 mRNA probe (green), lysozyme (pink), E-cadherin (red), DAPI (blue). DR3 expression on Paneth cells (arrowheads), and IECs (arrows). **(B-E)** Human intestinal organoids were cultured on a Paneth cell-Chip under continuous media flow with or without TL1A. **(B)** Expression of human DR3 mRNA on Paneth cell-Chip. DR3 (green), E-cadherin (red), DAPI (blue). DR3 expression on IECs (arrows). **(C)** Expression of human lysozyme (green), E-cadherin (red), and DAPI (blue). Top: Confocal microscopy imaging of untreated Paneth cell-Chip depicting Paneth cells with mature granule phenotype and predominant apical lysozyme expression (magnification: 63x (right), 40x (left)). Bottom: TL1A-treated Paneth cell-Chip depicting diffuse (D3), and disordered granules (D1) Paneth cell morphology. **(D)** Percentage of Paneth cells with D0, D1, D2, or D3 features. A total of 84 (control) and 113 (TL1A treated) Paneth cells were analyzed from 2 independent experiments. **(E)** Quantification of lysozyme immunoreactivity. Pooled data from 2 independent experiments. **(F-H)** Mouse ileal organoids were cultured under Paneth cell-enrichment conditions with or without TL1A. **(F)** Expression of lysozyme (red), E-cadherin (green), and DAPI (blue). Confocal microscopy imaging of control (left) or TL1A treated (right) Paneth cell-enriched organoids (magnification: 63x). White arrow heads indicate predominant apical lysozyme expression and white arrows indicate disordered lysozyme-containing granules. **(G)** Quantification of lysozyme fluorescence intensity (arbitrary units). A total of 224 (control) and 174 (TL1A treated) enteroids from 2 independent experiments were analyzed. **(H)** Numbers of Paneth cells per whole 3D reconstructed Paneth cell enriched enteroids. A total of 28 (control) and 32 (TL1A treated) enteroids from 3 independent experiments were analyzed. Data represent means ± SD. \**p*<0.05; \*\*\**p*<0.005, two-tailed Student’s *t*-test.

### TL1A alters the phenotype of Paneth cells in a human Paneth cell-Chip and Paneth cell enriched mouse enteroids

In order to study the direct effects of TL1A on human Paneth cells, we developed a human intestinal organoid-derived Paneth cell chip. We generated human intestinal organoids from healthy subjects and seeded a single-cell suspension on a micro-engineered Chip subjected to continuous laminar flow (33). To study Paneth cell phenotypes, we adapted our published culture protocol to enrich for Paneth cells by adding DAPT, a γ-secretase inhibitor, as described for biopsy-derived organoids (34). Consistent with human small intestinal Paneth cells, we observed DR3 expression in the Paneth cell-Chip (Figure 3B). We observed the formation of intestinal folds containing over 90% of lysozyme^+^ Paneth cells (Figure 3C). Furthermore, we observed a mature Paneth cell phenotype as characterized by predominately apical localization of lysozyme containing granules and secretion of lysozyme (Figure 3C and data not shown).

Compared to untreated Paneth cell-Chip, treatment with TL1A for four days resulted in an altered Paneth cell phenotype. We observed the majority of Paneth cells displayed an abnormal D1 and to a lesser degree D3 granule phenotype in TL1A treated Paneth cell-Chip (Figures 3C, D), and overall reduced intensity of lysozyme staining (Figure 3E). The D1, D3-dominant abnormalities in TL1A-treated Paneth cell-Chip is consistent with our findings of significant association with a D1, D3 Paneth cell phenotype and high mucosal TL1A expression in CD patients (Figure 1H, Table 1). Next, we enriched WT mouse enteroids for Paneth cells and treated them with TL1A. Consistent with our findings in human Paneth cells, we observed reduced intensity of lysozyme staining and predominantly disordered granule localization in the presence of TL1A, while control Paneth cell enriched enteroids were characterized by predominately apical localization of lysozyme containing granules (Figure 3F, G). We did not observe a change in the numbers of Paneth cells per enteroid upon treatment with TL1A or a change in mRNA expression of Paneth cell signature genes (Figure 3H, Supplemental Figure 1C).

### TL1A induces transcriptional changes in enteroids associated with mitochondrial function and autophagy

To determine direct effects of TL1A on downstream signaling pathways in intestinal epithelial cells, we treated ileal enteroids with TL1A and performed RNA-Sequencing analysis. We identified 19 genes differentially expressed in control vs TL1A treated enteroids at adjusted *p* < 0.05 (Figure 4A). Enrichment and Ingenuity Pathway analysis of differentially expressed transcripts revealed significant overrepresentation of biological processes of autophagy, mitochondrial membrane potential, and calcium import into mitochondria (Figure 4B). These results are consistent with mitochondrial abnormalities that we observed in *Tl1a-tg* mice (Figure 1G).

**Figure 4.**
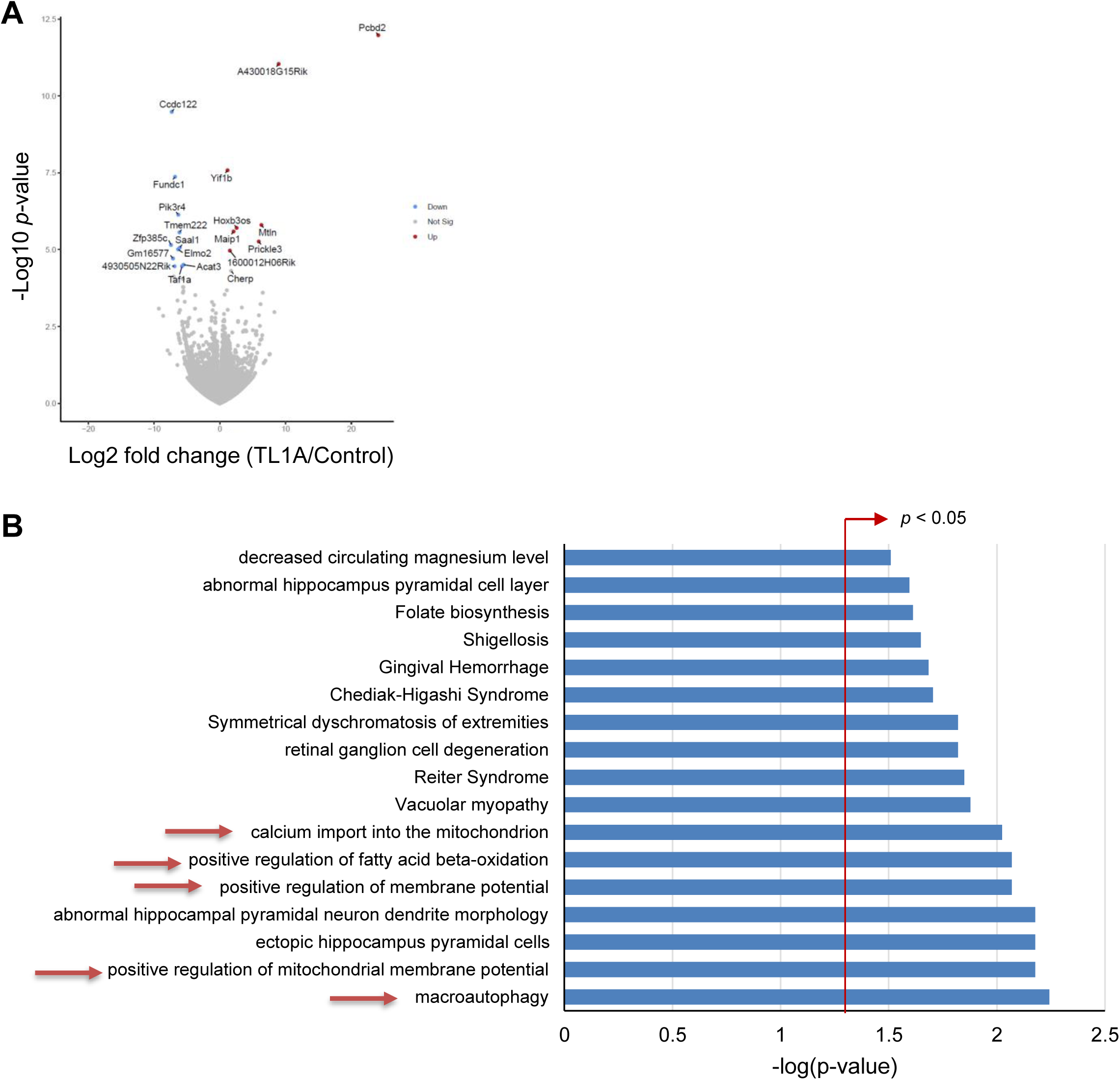
TL1A induces transcriptional changes in enteroids associated with mitochondrial function and autophagy. (**A**) Volcano plot showing significantly up and down-regulated genes in organoids treated with TL1A (n = 4 mice/group). (**B**) Pathway and Gene Ontology analysis of DEG in control vs. TL1A treated organoids.

### Intestinal microbiota is required for normal Paneth cell morphology and development of ileitis

As the microbiome is relevant to inflammation in numerous diseases including IBD, we evaluated the effects of GF conditions on TL1A-mediated intestinal inflammation and Paneth cell abnormalities. The absence of intestinal microbiota completely abrogated the spontaneous ileitis induced by TL1A overexpression, as there were no significant differences in ileal histopathology between GF *Tl1a-tg* and WT mice (Figure 5A). In *Tl1a-tg* mice we observed a significant increase in the numbers of Paneth cells in aged mice that we did not observe in WT mice under GF conditions (Figure 5B). However, we observed a lack of Paneth cell granule maturation in GF WT and *Tl1a-tg* mice at 5 and 10 months of age with a D3-dominant Paneth cell phenotype in both genotypes (Figures 5C, D). These data suggest that resident microbiota is required for Paneth cell granule maturation in WT and *Tl1a-tg* mice and the development of ileitis in *Tl1a-tg* mice, but dispensable for Paneth cell expansion in *Tl1a-tg* mice.

**Figure 5.**
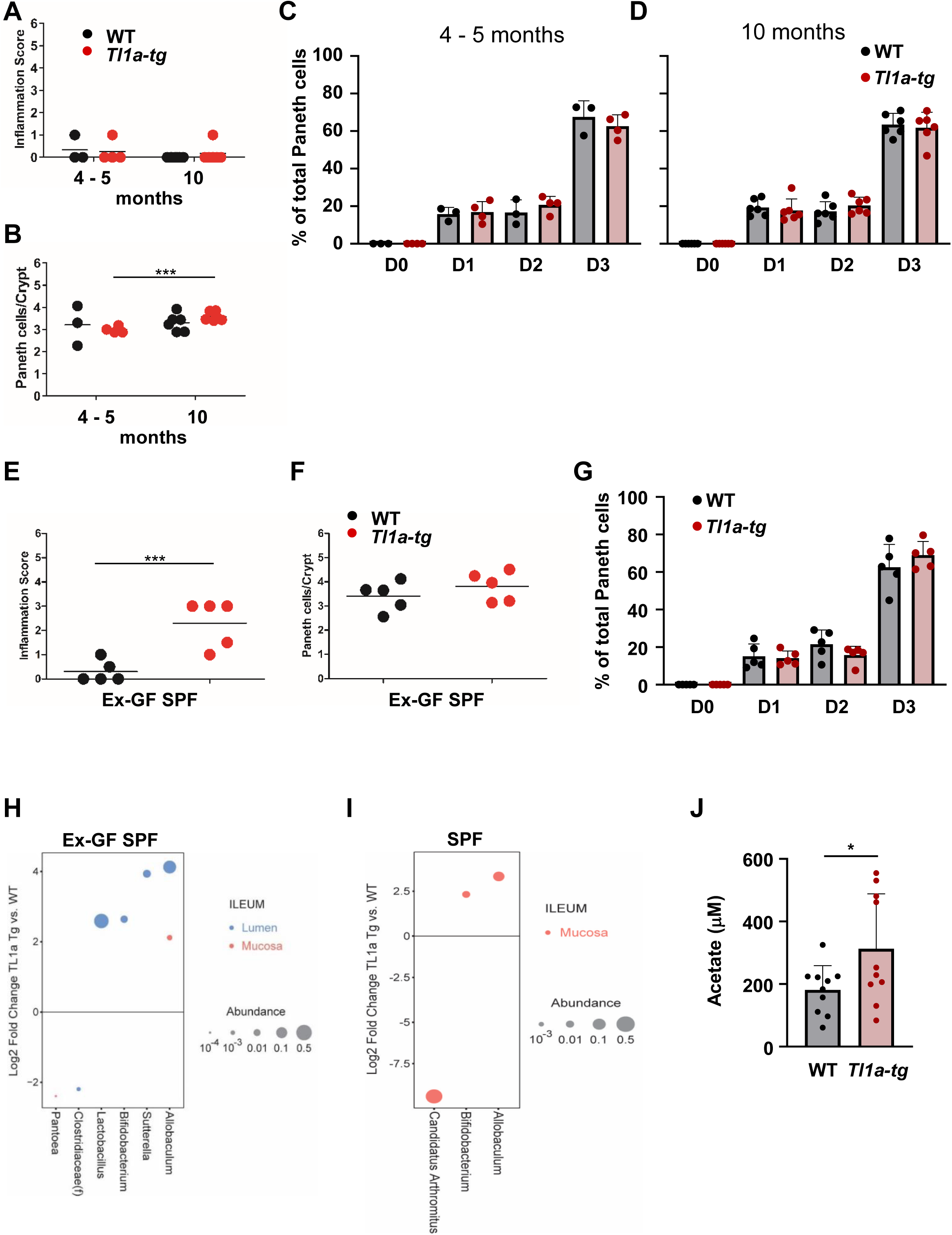
Intestinal microbiota are required for Paneth cell granule maturation and ileal inflammation. (**A - D**) Paneth cell phenotypes in germ-free (GF) WT and *Tl1a-tg* mice. (**A**) Inflammation score. (**B**) Numbers of Paneth cells. Percentage of Paneth cells with D0-3 features at 4-5 (**C**) and 10 months of age (**D**). Data represent means ± SD (n = 3-6 mice/group). (**E - G**) Paneth cell phenotypes in GF mice reconstituted with microbiota from SPF mice at 2-4 months of age. (**E**) Percentage of Paneth cells with D0-3 features at 10 months of age. (**F**) Numbers of Paneth cells. (**G**) Inflammation score (n = 3-6 mice/group). (**H**) GF WT or *Tl1a-tg* mice were reconstituted with SPF microbiota. 16s rRNA sequencing was performed of luminal and mucosal ileal samples from littermate controls 2 months after reconstitution (n = 10/group). Differential microbial genera (or unclassified members of Clostridiaceae) were identified by negative binomial models. Log2 of the fold change between *Tl1a-tg* and WT is shown. Dot size represents microbial relative abundance and color denotes sample type (lumen vs. mucosa). (**I**) 16s rRNA sequencing was performed of luminal and mucosal ileal samples from 2-month-old SPF WT or *Tl1a-tg* mice littermate controls (n = 10/group). Log2 of the fold change between *Tl1a-tg* and WT is shown. (**J**) Measurement of short chain fatty acids in ileal content from SPF littermate controls (n = 10/group). Acetate was more abundant in the ileum of SPF *Tl1a-tg* mice. \**p*<0.05; \*\*\**p*<0.005 by two-tailed Student’s *t*-test.

### SPF microbiota promotes intestinal inflammation and Paneth cell expansion

Next, we reconstituted GF mice with murine SPF microbiota to determine if microbiota reconstitution can restore the development of ileitis and Paneth cell morphology. GF mice were reconstituted with SPF microbiota (Ex-GF SPF) at 2 months of age by oral gavage and inflammation and Paneth cell morphology was analyzed at 5 months of age. Reconstitution with SPF microbiota restored the development of ileitis in *Tl1a-tg* mice (Figure 5E). However, SPF reconstitution did not restore Paneth cell expansion or led to Paneth cell granule maturation (Figures 5F, G). Our data indicate that reconstitution of adult GF mice is sufficient to restore the development of ileitis in *Tl1a-tg* mice but insufficient for Paneth cell expansion and granule maturation.

### Abnormalities in Paneth cell morphology is associated with changes in microbiota and the metabolite acetate in Tl1a-tg mice

Next, we performed 16S rRNA sequencing to characterize the ileal microbiome of colonized ex-GF and SPF mice. Reconstitution of adult GF mice with WT SPF feces was associated with higher abundance of luminal *Lactobacillus, Bifidobacterium, Sutterella,* and luminal and mucosa-associated *Allobaculum* in ex-GF *Tl1a-tg* compared to ex-GF WT mice suggesting that these bacterial strains might drive development of ileitis (Figure 5H). Next, we analyzed the abundance of microbial genera under SPF conditions in 2-month-old WT and *Tl1a-tg* mice. We observed a higher abundance of ileal, mucosal-associated *Bifidobacterium,* and *Allobaculum* in *Tl1a-tg* mice (Figure 5I). Next, we performed targeted metabolic profiling of ileal contents of 2-month-old SPF WT and *Tl1a-tg* mice. We observed a significant increase in the concentration of the short-chain fatty acid acetate in *Tl1a-tg* mice while concentrations of butyrate and propionate were similar between WT and in *Tl1a-tg* mice (Figures 5J, Supplemental Figure 2). The observed increase in acetate is consistent with the increase of the abundance of the acetate producers *Bifidobacterium* and *Allobaculum* in *Tl1a-tg* mice.

### Short-chain fatty acid acetate drives ileal inflammation

Next, we examined the impact of acetate on the development of ileitis. We first treated 12-week-old littermate GF WT or GF *Tl1a-tg* mice with 200 mM sodium acetate in drinking water (*ad libitum*) for 3 months (Figure 6A). Both, GF WT and GF *Tl1a-tg* mice supplemented with acetate developed ileal inflammation compared to mice receiving regular drinking water (Figures 6B, C). However, acetate supplementation did not result in normal Paneth cell granule maturation independently of the genotype of the mice suggesting that acetate supplementation alone is not sufficient to drive normal Paneth cell granule development (Figure 6D). We also did not observe any significant differences in the numbers of Paneth cells/crypt in acetate treated GF WT and GF *Tl1a-tg* mice (Figure 6E). However, we observed significantly elevated IFN-γ secretion by MLN in acetate treated WT and *Tl1a-tg* mice (Figure 6F). In contrast, we did not observe any significant increase in intestinal permeability in acetate treated mice (Figure 6G). Consistent with previous publications, we observed a significant increase of the numbers of MLN Treg populations in WT mice receiving acetate supplementation (Supplemental Figure 3A) (35, 36). We observed a significant increase in the percentage of FoxP3^+^ RORγt^+^ Tregs and particularly in the percentage of proliferating FoxP3^+^ RORγt^+^ Tregs in WT and *Tl1a-tg* mice upon acetate supplementation (Supplemental Figure 3B). Our data demonstrate that acetate is sufficient to drive ileitis in the absence of microbiota independently of the host genotype. In a second set of experiments, we treated SPF WT or *Tl1a-tg* mice with acetate in drinking water for 2 or 4 months starting at 2 months of age (Figure 7A). WT mice treated with acetate for 4 months developed ileal inflammation compared to water controls (Figure 7B, D). In *Tl1a-tg* mice acetate treatment for 2 months led to ileitis that was not observed in age-matched water control *Tl1a-tg* mice suggesting an acceleration of the development of ileal inflammation in *Tl1a-tg* mice by acetate (Figure 7B, C). In addition, we observed significantly reduce percentages and total numbers of MLN naïve T cells and concomitant increased percentages and numbers of effector CD4^+^ T cells in acetate treated WT mice consistent with our findings of ileal inflammation in these mice (Supplemental Figure 4A, B).

**Figure 6.**
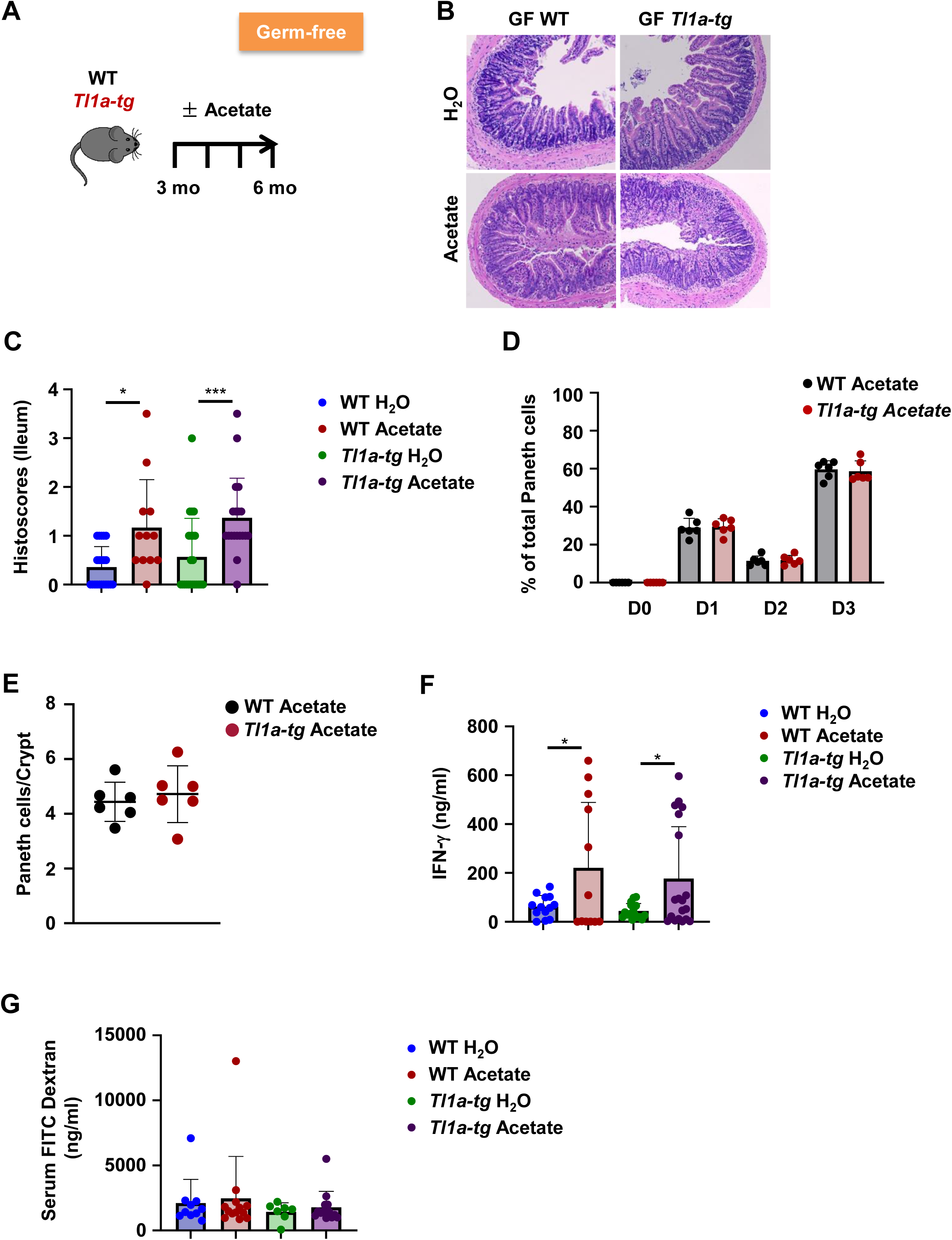
The microbial metabolite acetate promotes ileal inflammation in GF mice. **(A)** Experimental setup for acetate supplementation of GF-free mice. **(B)** Histoscores (n = 12-22/group). **(C)** Representative H & E staining in water or acetate treated GF WT and *Tl1a-tg* mice. **(D)** Percentage of Paneth cells with D0-3 features of water or acetate treated GF WT or *Tl1a-tg* mice (n = 6/group). **(E)** Numbers of Paneth cells/crypt (n = 6/group). **(F)** Anti-CD 3 and anti-CD28 antibody-stimulated IFN-γ secretion of MLN of water or acetate treated GF WT or *Tl1a-tg* mice (n = 12-19/group). **(G)** FITC dextran serum concentration (n = 7-13/group). \**p*<0.05, ** *p*<0.01 by Mann-Whitney U test (B) or Student’s *t*-test.

**Figure 7.**
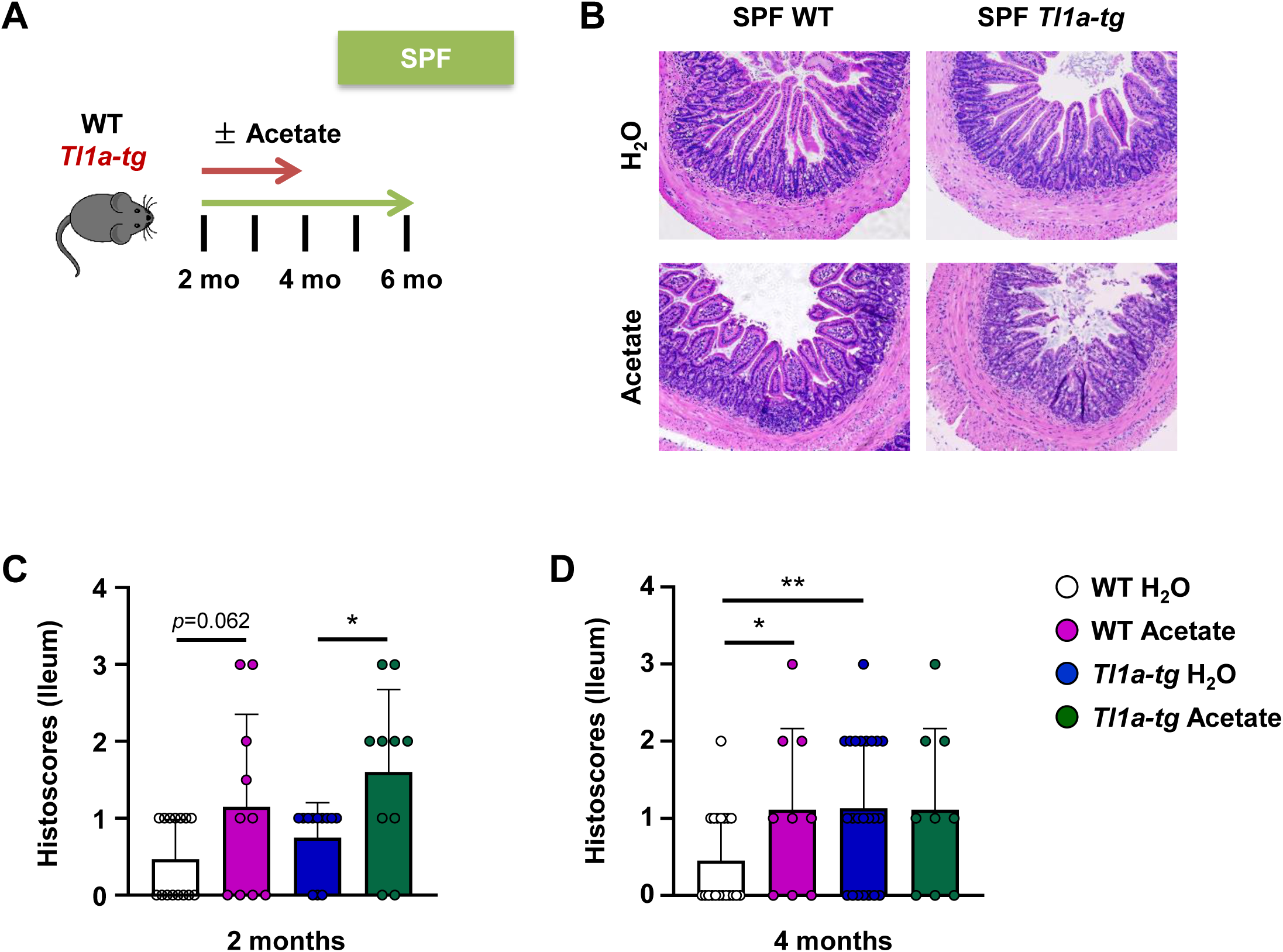
The microbial metabolite acetate promotes ileal inflammation in SPF mice. **(A)** Experimental setup for acetate supplementation of SPF mice. **(B)** Representative H & E staining in water or acetate treated SPF WT and *Tl1a-tg* mice. **(C, D)** Histoscores after 2 months **(C)** and 4 months **(D)** treatment (n = 9-23/group). \**p*<0.05, ** *p*<0.01 by Student’s *t*-test.

### Paneth cell-intrinsic DR3 signaling pathway has a direct impact on Paneth cell morphology

Based on our observation that DR3 is expressed on Paneth cells, we hypothesized that TL1A might directly affect Paneth cell function. We generated mice with specific deletion of DR3 in Paneth cells (*Dr3^ΔPC^*) by crossing floxed DR3 mice with the Defa6-Cre line (13). Next, we crossed *Dr3^ΔPC^*mice to *Tl1a-tg* mice (*Tl1a-tg Dr3^ΔPC^* mice) to assess direct and indirect effects of TL1A on Paneth cell morphology and function. In comparison to WT mice, *Dr3^ΔPC^* and *Tl1a-tg Dr3^ΔPC^* mice displayed a significantly higher degree of Paneth cell abnormalities at any age but most pronounced in older mice and these abnormalities preceded the development of ileitis (Figures 8A, D). Compared to *Tl1a-tg Dr3^ΔPC^* mice, Paneth cell abnormalities are less severe in *Dr3^ΔPC^* mice at 6 months of age suggesting that direct and indirect effects of TL1A overexpression on Paneth cells contribute to the abnormal morphology (Figure 8D). In addition, similar Paneth cell numbers were observed in *Dr3^ΔPC^* mice compared to WT in 2 – 4, and 6-month-old mice (Figures 8B, E). In contrast, *Tl1a-tg Dr3^ΔPC^* mice have significantly increased Paneth cell numbers compared to WT and *Dr3^ΔPC^* mice suggesting that Paneth cell hyperplasia is driven by TL1A overexpression in a manner independent of DR3 expression on Paneth cells. Spontaneous ileitis was not observed in *Dr3^ΔPC^* mice but in 6-month-old *Tl1a-tg Dr3^ΔPC^* mice (Figures 8C, F). Compared to *Tl1a-tg* mice, the ileitis observed in 6-month-old *Tl1a-tg Dr3^ΔPC^*mice is less severe suggesting that the TL1A-DR3 pathway contributes to ileitis in a Paneth cell-intrinsic and -extrinsic manner (Figure 8F). Therefore, DR3 signaling on Paneth cells contributes to the maintenance of normal granule packaging in Paneth cells. Next, we performed RNA-Sequencing analysis on ileal crypts from 3-months old *Dr3^fl/fl^*, *Dr3^ΔPC^*, and *Tl1a-tg Dr3^ΔPC^* mice. In *Tl1a-tg Dr3^ΔPC^* ileal crypts, we observed transcriptional changes in pathways associated with the unfolded protein response and ER protein processing compared to *Dr3^ΔPC^* crypts and pathways involved in protein localization and mitochondria morphogenesis compared to *Dr3^fl/fl^* crypts. In *Dr3^ΔPC^* ileal crypts, we observed transcriptional changes in pathways associated with cellular metabolism compared to *Dr3^fl/fl^* crypts (Supplemental Figures 5, 6). Next, we determined if there are any changes in the LP and MLN immune cell compositions in *Dr3^ΔPC^*, and *Tl1a-tg Dr3^ΔPC^* mice. We did not observe any significant differences in LP and MLN total cell numbers, naïve and effector CD4^+^ T cell percentages or total numbers under homeostatic conditions in *Dr3^ΔPC^*, and *Tl1a-tg Dr3^ΔPC^* mice (Supplemental Figures 7<u>, 8</u>). However, we observed a significant increase in the percentage of FoxP3^+^ RORγt^+^ Tregs and particularly in the percentage of proliferating FoxP3^+^ RORγt^+^ Tregs in *Tl1a-tg Dr3^ΔPC^* mice (Supplemental Figures 7).

**Figure 8.**
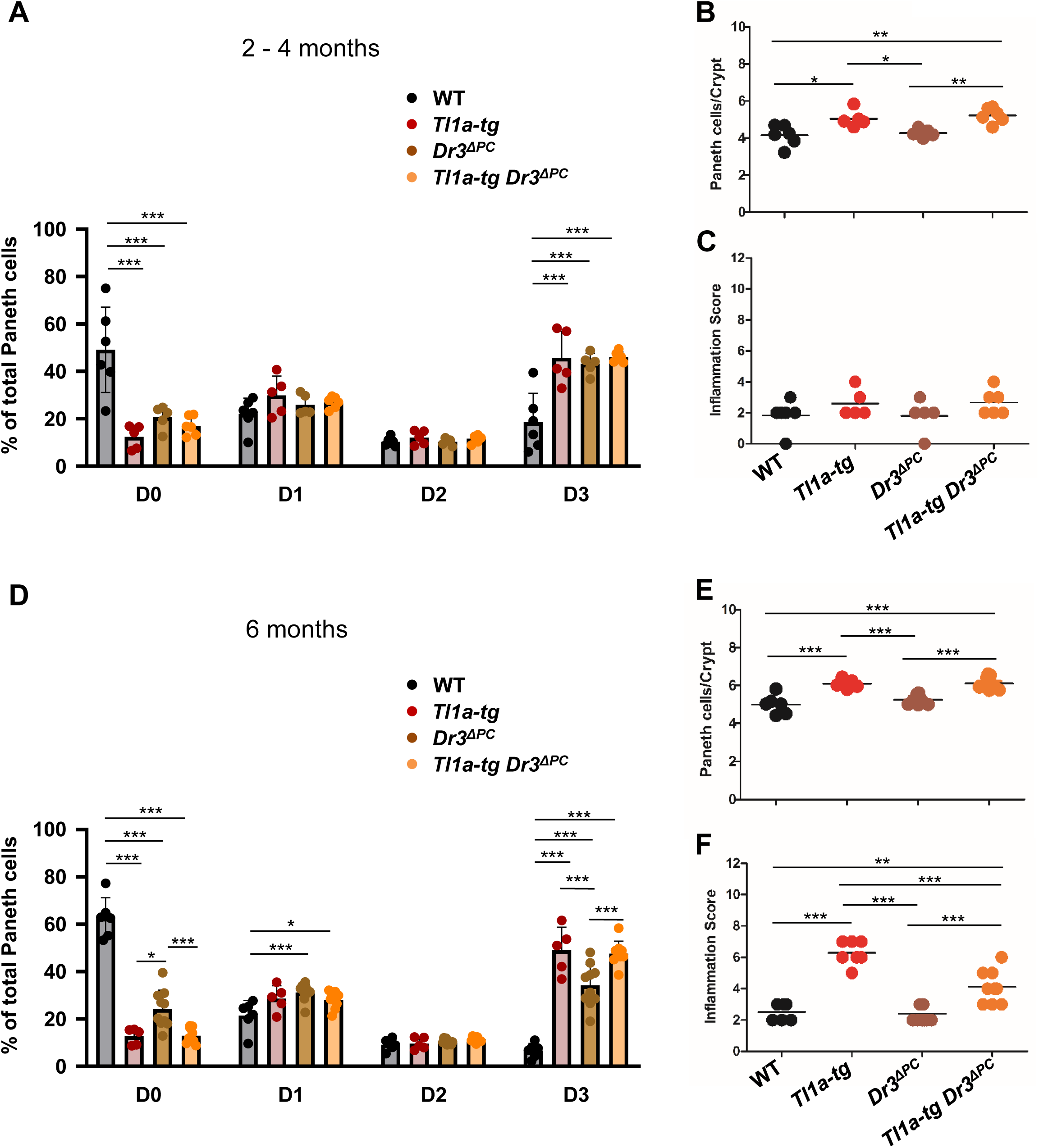
DR3 signaling regulates development and function of ileal Paneth cells in a cell-intrinsic manner. (**A-C**) Paneth cells Phenotype in *WT, Tl1a-tg, Dr3^ΔPC^*, and *Tl1a-tg Dr3^ΔPC^* mice at 2 – 4 months of age. Data represent means ± SD (n = 5-10 mice/group). (**A**) Percentage of Paneth cells with D0-3 features. (**B**) Numbers of Paneth cells. (**C**) Inflammation score. (**D-F**) Paneth cells Phenotype in *WT, Tl1a-tg, Dr3^ΔPC^*, and *Tl1a-tg Dr3^ΔPC^* mice at 6 months of age. Data shown as means ± SD (n = 5-10 mice/group). (**D**) Percentage of Paneth cells with D0-3 features. (**E**) Numbers of Paneth cells. (**F**) Inflammation score. \**p* < 0.05; \*\**p* < 0.01; \*\*\**p* < 0.005, 1-way ANOVA with Tukey’s HSD test for more than two groups.

### Defects in Paneth cell-intrinsic DR3 signaling pathway is associated with changes in microbiota composition

Next, we performed 16S rRNA sequencing to characterize the ileal microbiome of *Dr3^ΔPC^*, and *Tl1a-tg Dr3^ΔPC^* mice. While we did not observe significant differences in alpha diversity between genotypes, PERMANOVA-based analysis of distance matrix indicated that the beta diversity of microbial communities was significantly different between *Dr3^fl/fl^*, *Dr3^ΔPC^*, and *Tl1a-tg Dr3^ΔPC^* mice (Figure 9A, *p* < 0.05). The microbial composition was relatively more diverse at the phylum, families, and genus level in *Dr3^ΔPC^*, and *Tl1a-tg Dr3^ΔPC^* mice when considering *Dr3^fl/fl^*mice as a reference group (Figure 9B, and data not shown). MaAslin2 analysis was used to identify differential microbial features in *Dr3^ΔPC^*, and *Tl1a-tg Dr3^ΔPC^* mice when considering *Dr3^fl/fl^* mice as a reference group. A total of 24 significantly abundant taxa were detected. Higher abundance of *Streptococcus*, *Ligilactobacillus* was observed in *Dr3^fl/fl^* mice, while *Dr3^ΔPC^*, and *Tl1a-tg Dr3^ΔPC^* mice had a higher abundance of *Lactobacillus*, *HT002*, and *Desulfovibrio* (Figures 9B-D). In addition, genus *Enterorhabdus* were specifically abundant in *Tl1a-tg Dr3^ΔPC^* mice.

**Figure 9.**
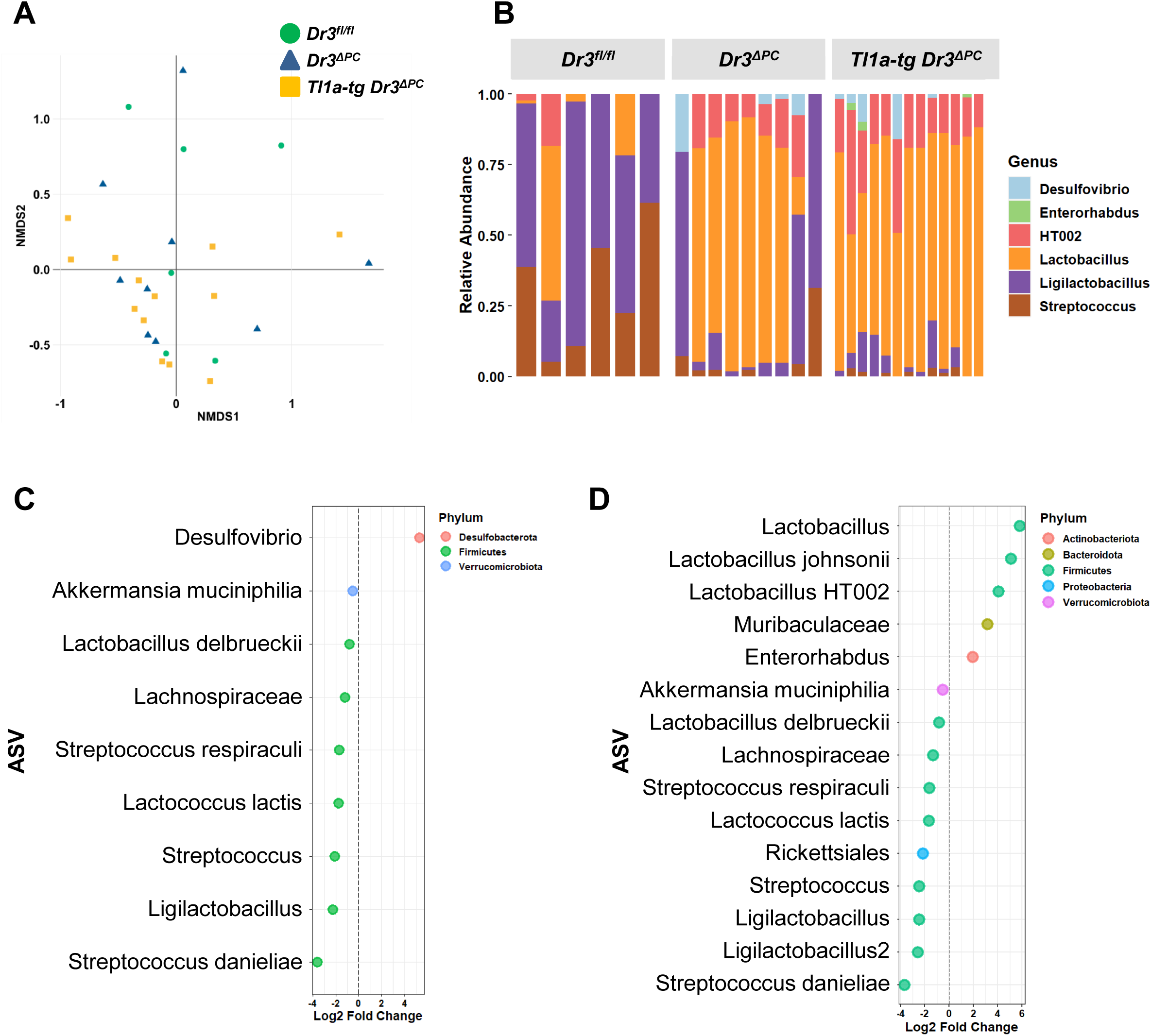
Effects of Paneth cell-intrinsic DR3 deletion on the ileal microbiota. (**A-D**) 16s rRNA sequencing was performed of luminal ileal samples from littermate *Dr3^fl/fl^*, *Dr3^ΔPC^*, and *Tl1a-tg Dr3^ΔPC^* mice at 2 months of age (n = 6-13/group). (**A**) Non-metric multidimensional scaling (NMDS) plot of 16s rRNA sequencing data based on PERMANOVA analysis (p < 0.05). (**B**) Relative abundance of ileal microbiota of *Dr3^fl/fl^*, *Dr3^ΔPC^*, and *Tl1a-tg Dr3^ΔPC^* mice. (**C**) Significantly differential abundant taxa were identified by negative binomial models. Log2 of the fold change between *Dr3^ΔPC^*and *Dr3^fl/fl^* is shown. Color denotes different phyla. (**D**) Log2 of the fold change between *Tl1a-tg Dr3^ΔPC^* and *Dr3^fl/fl^* is shown. Color denotes different phyla.

## Discussion

The interplay between intestinal microbiota and their metabolites and the intestinal epithelial barrier, and ensuing host immune responses is important in the etiology of human IBD. Our data highlight a role of the IBD-associated genes *TNFSF15* (encoding TL1A) and its receptor *TNFRSF25* (encoding DR3) in Paneth cell dysfunction leading to the development of ileitis. We report a significant correlation between levels of ileal *Tl1a* mRNA expression in CD patients and Paneth cell abnormalities independently of the *ATG16L1* genotype. Furthermore, we did not observe a significant association between the expression of the IBD-associated gene *CARD15* (encoding NOD2) and Paneth cell abnormalities in our dataset which may be related to the uninflamed ileal tissue sample that we analyzed. Using a highly specific and sensitive probe for *Dr3*, we detected *Dr3* mRNA expression on murine Paneth cells by smFISH. Moreover, we also observed *Dr3* expression on human Paneth cells in surgical mucosal sections of CD patients and on human iPSC derived human intestinal organoids. Mice with a deletion of DR3 on Paneth cells (*Dr3^ΔPC^*) showed a predominantly abnormal Paneth cell phenotype while mice with a deletion of DR3 on Paneth cells and TL1A overexpression (*Tl1a-tg Dr3^ΔPC^*) showed a more severe abnormal Paneth cell phenotype. Our data suggest that Paneth cell intrinsic and extrinsic TL1A-DR3 signaling contribute to abnormal Paneth cell morphology potentially through the induction of IFN-γ in *Tl1a-tg* mice. A direct effect of TL1A on Paneth cell morphology was confirmed in human *in vitro* Paneth cell Chip cultures. TL1A treatment resulted in abnormal Paneth cell phenotype with the majority of Paneth cells displaying abnormal D1 and to a lesser degree D3 granule phenotype, and overall reduced intensity of lysozyme staining. The D1, D3-dominant abnormalities in the TL1A-treated Paneth cell Chip are consistent with our findings of significant association with a D1, D3 Paneth cell phenotype and high mucosal TL1A expression in CD patients. Our data demonstrate the usefulness of our Paneth cell Chip in modeling human Paneth cell morphology and function to elucidate signaling pathways leading to Paneth cell dysfunction in CD patients. In addition to the direct effect TL1A has on Paneth cells, our data also demonstrate that indirect effects, most likely via the pro-inflammatory effects TL1A exerts on effector T cells, contribute to Paneth cell abnormalities and the development of ileitis.

Interestingly, TL1A overexpression results in Paneth cell hyperplasia rather than Paneth cell loss, which is usually observed in patients with CD and in mouse models of Paneth cell dysfunction (37). This suggests that TL1A has diverse effects on Paneth cell biology including hyperplasia and granule morphology. The Paneth cell hyperplasia that we observed in mice overexpressing TL1A is independent of DR3 expression on Paneth cells. IL-9 has been shown to drive IL-13 dependent hyperplasia (38). We previously showed that TL1A upregulated IL-9 in T helper cells and IL-13 overexpression has also been shown in *Tl1a-tg* mice (8, 39). Future studies investigating a potential role of Paneth or epithelial cell-intrinsic TL1A on Paneth cell phenotypes and function will be required to dissect the potential role of epithelial-intrinsic TL1A from the role of exogenous TL1A presented in this study.

IBD results from dysregulated immune responses to intestinal microbiota in a susceptible host. Previous studies have reported major shifts in the gut microbiota composition in patients with IBD (40, 41). However, a recent large scale longitudinal study in patients with IBD attributed the majority of variance to inter-individual variations rather than disease status and reported longitudinally changes correlating with disease activity states (18). Furthermore, they identified metabolome changes as the most apparent changes amongst the parameters analyzed (compared to bacterial metagenomics, metatranscriptomics) (18). Amongst the changes in metabolomic profiles in IBD patients a reduction of short chain fatty acids (SCFA) and particularly butyrate and propionate and increase in primary bile acids and sphingolipids could be observed particularly in IBD patients with dysbiosis (18, 20). However, one of the limitations of both studies is the reliance of stools samples for analysis, which might be less representative for metabolic profiles in the ileum. Interestingly, it has been proposed that Paneth cell defects may induce dysbiosis, but it is still controversial whether dysbiosis is a consequence or instigator of Paneth cell dysfunction since both profound changes in microbiota and metabolites and Paneth cell abnormalities precedes the development of ileitis (13, 42, 43). Our studies support a direct role of resident microbiota, with the absence of ileitis in GF *Tl1a-tg* mice and restored inflammatory phenotype following SPF microbial restitution in adult ex-GF *Tl1a-tg* mice.

Consistent with these findings, we observed significant Paneth cell abnormalities and changes in microbiota composition in *Tl1a-tg* mice to precede ileitis in these mice. Furthermore, *Tl1a-tg* mice were enriched in acetate-producing bacteria *Bifidobacterium* and *Allobaculum* and metabolomic analysis revealed an increase of the metabolite acetate in the ileum of *Tl1a-tg* mice. The short chain fatty acid acetate has been mainly associated with anti-inflammatory effects including its positive effects on the suppressive function of Tregs (44). However, acetate has also been shown to promote effector T cell function and particularly TH1 and TH17 responses (45). Our data are consistent with these findings, and we report a significant increase in IFN-γ production in MLN of acetate-treated GF mice independent of the genotype of the mice which correlated with the development of ileitis. IFN-γ has been reported to induce apoptosis and loss of Paneth cells in the context of infection with the protozoan parasite Toxoplasma gondii or T cell activation dependent inflammation (28, 46) suggesting that IFN-γ could further exacerbate Paneth cell dysfunction.

Interestingly, we observed significantly more diverse ileal microbial communities in both, *Dr3^ΔPC^* and *Tl1a-tg Dr3^ΔPC^* mice compared to *Dr3^fl/fl^* mice. *Dr3^ΔPC^*, and *Tl1a-tg Dr3^ΔPC^* mice had a higher abundance of *Lactobacillus*, *HT002*, and *Desulfovibrio,* the abundance of the latter being associated with IBD (47).

Our data also suggest that a tight regulation of TL1A-DR3 signaling on Paneth cells is required for normal Paneth cell morphology and microbial composition.

In conclusion, our study provides a mechanistic link between TL1A overexpression, Paneth cell dysfunction, and intestinal microbiota, all of which have been shown to contribute to IBD. Overexpression of TL1A leads to alterations in Paneth cell numbers, morphology and function that promote ileal inflammation in a microbiota-dependent manner via the microbial metabolite acetate. Acetate contributes to the development of ileitis by enhancing TH1 responses. Our data have implications for how microbiota-derived acetate may elicit previously unknown pro-inflammatory effects in the context of genetic IBD risk factors.

## Methods

### Sex as a biological variable

Our study examined male and female animals, and similar findings are reported for both sexes.

### Mice

Age- and sex-matched mice on the C57BL/6J background were used. Mice overexpressing TL1A in CD4^+^ T cells (LCK-CD2-Tl1a-GFP; *Tl1a-tg*)(7) and wild-type (WT) littermate control mice were used. Mice containing a floxed Dr3 allele (*Dr3^fl/fl^*) were generated by GenOway as described (48). Defa6-Cre mice were kindly provided by Richard Blumberg (Brigham and Women’s Hospital, Boston, MA) (13). Defa6-Cre were crossed to *Dr3^flox/flox^* to generate Paneth-cell-specific deletion of DR3 (*Dr3^ΔPC^*). *Tl1a-tg* mice were crossed to *Dr3^ΔPC^* to generate *Tl1a-tg Dr3^ΔPC^*. Mice were maintained under specific-pathogen-free (SPF) or germ-free (GF) conditions.

### GF and microbiota reconstituted mice

*Tl1a-tg* and WT mice were re-derived into GF status at the National Gnotobiotic Rodent Resource Center (University of North Carolina, Chapel Hill). 2-4 months old *Tl1a-tg* mice and WT littermates were orally gavaged with 200 µl of a 1:10 suspension of stool collected from WT Cedars-Sinai specific pathogen free (SPF) mice diluted in phosphate-buffered saline as described (48). GF mice were euthanized at 4-5 months or 10 months of age and reconstituted mice were euthanized at 5 months of age.

### Acetate supplementation

Sodium acetate (Sigma-Aldrich) was added at 200 mM to the drinking water. SPF mice were supplemented with acetate water for 2 or 4 months starting at 2 months of age. GF mice were supplemented with acetate water for 3 months starting at 3 months of age. Mice had ad libitum access to water and food during the entire treatment period.

### Intestinal Permeability Assay

Mice were fasted overnight and orally gavaged with FITC-dextran (molecular weight, 3000–5000 daltons, Sigma; 500 mg/kg body weight). Blood was collected by retro-orbital bleeding 1 hour after gavage and serum was obtained and stored at -80°C until use. The serum FITC-dextran concentration was measured with a fluorescence spectrophotometer (SpectraMax; Molecular Devices) at an excitation/emission of 485 nm/520 nm and analyzed using a standard curve of FITC dextran diluted in PBS. Mouse serum from untreated mice was used as a background control.

### Histology Scoring

1-2 cm of ileal tissues were collected near the cecum, paraffin-embedded and stained with H&E for histological analysis. Severity of ileal inflammation was evaluated in a blinded manner according to previously described criteria by two trained animal pathologists. Spontaneous ileitis was evaluated by scoring the degree of inflammation, extent of involvement, crypt damage and villus changes as described (49). GF mice were evaluated using the scoring criteria described by Erben et al. (50).

### 16S rRNA gene amplicon sequencing

Ileal luminal content was released by flushing with distilled deionized water then the mucosa-associated bacteria were released by DTT treatment according to published protocols (51). DNA extraction and sequencing of the 16S ribosomal RNA gene was then performed for luminal and mucosal samples as previously described (51). In brief, bacterial DNA was extracted using the DNeasy PowerSoil Pro kit (Qiagen) with bead beating according to the manufacturer’s instructions. The V4 region of the 16S rRNA gene was amplified using barcoded primers containing Illumina flow-cell compatible adapter sequences. The resulting amplicons were pooled in equal amounts to create sequencing libraries, which were sequenced on Illumina MiSeq (2 x 250 bp) or on Illumina HiSeq 2500 (2 x 150 or 2 x 250 bp).

### Bioinformatics analysis of the 16S amplicon data

For microbiome analysis of WT and *Tl1a-tg* ileal luminal and mucosal samples, raw data was processed in QIIME 1.9.1 and 97% operational taxonomic units (OTUs) were identified by closed reference OTU picking against the Greengenes database. Differentially abundant genera were identified by negative binomial models implemented in DESeq2 (51). For microbiome analysis of *Dr3^fl/fl^*, *Dr3^ΔPC^*, and *Tl1a-tg Dr3^ΔPC^* ileal content, demultiplexed sequence data was analyzed using Divisive Amplicon Denoising Algorithm 2 (DADA2), version 1.16 pipeline. Briefly, forward and reverse reads were filtered and trimmed based on the sequence quality assessment. Next, for forward and reverse reads, a model was built to learn the error rate, and the forward and reverse reads were dereplicated to combine all identical sequencing reads into unique sequences. Finally, error model and dereplicated sequence data were used to infer Amplicon Sequence Variants (ASVs) using the DADA2’s core denoising algorithm – dada and ASV table was built after merging forward and reserves reads. Next the chimeric sequences were removed using consensus method. Taxonomy was assigned to ASV using Silva database (version 138.1). A phyloseq object was created using ASV table and taxonomic table, and subsequent diversity analyses was performed in the phyloseq R package (version 1.46). Alpha diversity was estimated after rarefying the data to the lowest sequence count of 8,233 sequences per sample. Four different alpha diversity indices (Shannon. Simpson, Observed ASVs, and Inverse Simpson) were calculated and tested for group differences using ANOVA test with post hoc analysis using Tukey’s HSD test. The Beta diversity analysis was conducted utilizing the Bray-Curtis dissimilarity matrix computed on the raw sequence counts. The relationships among samples in relation to their group affiliations were visually represented through a Non-Metric Multidimensional Scaling (NMDS) plot. Next, the significance of group affiliation on beta diversity was tested using permutational multivariate analysis of variance (PERMANOVA). Finally, differentially abundant ASVs were estimated using MaAsLin2 package. For MaAsLin2 analysis, normalization method used was TMM and the data was log transformed before estimating the differentially abundant taxa with a minimum prevalence of 10% samples for which a ASV is detected. 16S rRNA sequences generated in this study are publicly available (NCBI BioProject: PRJNA951424).

### Metabolomic analysis

For metabolomics analysis, the aqueous phase aliquots of homogenized luminal fecal samples in ddH2O were sent to the West Coast Metabolomics Center, a NIH-supported core service located at UC Davis. Targeted gas chromatography quadrupole mass spectrometry analyses were performed to measure short chain fatty acids.

### Immunofluorescence staining of lysozyme

Lysozyme staining of ileal tissue sections was performed as described (22). In brief, formalin-fixed, paraffin-embedded tissue sections were deparaffinized, blocked with hydrogen peroxide, and boiled in citrate buffer (10 mM Sodium Citrate, 0.05% Tween-20; pH 6.0) for antigen-retrieval. Tissue sections were stained overnight at 4° C with a goat polyclonal anti-Lysozyme antibody (Santa Cruz Biotechnology, CA; sc-27958) at a 1:150 dilution and followed by secondary antibody Alexa Fluor^®^ 594-conjugated donkey anti-goat IgG (Molecular Probes, A-11058) for 2 h at room temperature (1:500 dilution). Total numbers of Paneth cells and percentages of Paneth cells with D0-D3 features in ileum were quantitated by an investigator blinded to the mouse genotypes as described (22). For each mouse, Paneth cells were analyzed in well-oriented ileal crypts and the percentages of D0, D1, D2, D3 Paneth cells were determined as percentage of total numbers of Paneth cells. Paneth cells were counted to determine the average number of Paneth cells per crypt in each mouse. For lysozyme staining of Paneth cell-Chips, chips were flushed through the upper and lower channels with PBS, fixed with 4% paraformaldehyde, incubated in 30% sucrose overnight at 4°C, and embedded in Tissue-Tek O.C.T Compound (VWR). Cross-sections of the chips were obtained using a Leica CM3050S cryostat. Sections were blocked in 10% normal donkey serum and incubated with anti-E-cadherin (AF648, R&D Systems) and anti-Lysozyme antibodies (NB100-63062, Novus Biologicals) for 24 h at 4°C, followed by secondary antibodies as describe above and DAPI counterstains. Images were captured with a Leica TCS spectral microscope and Leica STELLARIS 8 STED and analyzed using Fiji Image J software. For lysozyme staining of murine organoids, whole mount enteroid staining was performed as described (52). Briefly, organoids were harvested using gentle cell dissociation reagent (STEMCELL Technologies), resuspended in 4% paraformaldehyde and incubated at 4°C for 45 minutes. Organoids were washed with PBS/0.1% Tween, blocked with PBS/0.1% Triton-x-100/0.2% BSA, and incubated with rabbit anti-Lysozyme (ab108508, Abcam, 1:100) and goat anti-E-cadherin (AF648, R & D Systems, 1:200) antibodies overnight at 4°C, followed by secondary antibodies donkey anti-goat AF 594 (A11058, Life technology, 1:500), and donkey anti-rabbit Dylight 650 (ab96922, Abcam, 1:500) overnight at 4°C, followed by counterstaining with Hoechst dye (62249, ThermoFisher Scientific).

### Single molecule Fluorescent in situ hybridization (smFISH)

Tissues were prepared as described for Immunofluorescence stainings. The DR3 transcripts were stained with RNAscope Multiplex Fluorescent Reagent Kit v2 (ACD, Newark, CA) according to manufacturer’s instruction followed by Immunofluorescence stainings for E-cadherin and lysozyme. The probe for murine *Dr3* transcripts was custom designed to hybridize between exons 2 and 5 of the murine *Dr3* which is deleted in the process of the generation of *Dr3*^-/-^ mice. The probe for human Dr3 was purchased from Advanced Cell Diagnostics (Catalog number: 448571). Probe for PPIB (Cyclophilin B) and Hela cell pellets were used as a positive control in this assay and a probe for DapB was used as a negative control.

### Laser Capture Microdissection for intestinal crypts, isolation of ileal crypts, RNA-sequencing, and data analysis

Ileal tissue sections were fixed in Methacarn (Methanol-Carnoy) fixation solution (60% methanol, 30% chloroform, 10% glacial acetic acid) for 1 h at room temperature. To compare the expression profile of Paneth cells between WT and *Tl1a-tg* mice, 5 cells at the bottom of the well-oriented crypts were obtained through Leica LMD7000 Laser Microdissection Cell Capture System. RNA from *Tl1a-tg* and WT crypts (n = 3/genotype) was extracted using Arcturus PicoPure® RNA Isolation Kits (ThermoFisher Scientific, Waltham, MA). Ileal crypts were isolated from 3 months old *Dr3^fl/fl^*, *Dr3^ΔPC^*, and *Tl1a-tg Dr3^ΔPC^* mice as described (53, 54). RNA was extracted using RNeasy Mini or RNeasy Micro kits (Qiagen). RNA-Seq libraries were created using the Kapa stranded mRNA-seq kit (Roche, Indianapolis, IN). Samples were sequenced using HiSeq 3000 (1 x 50 nt) or NovaSeq 6000 (for isolated crypts and organoids; 2 x 50 nt) (Illumina, San Diego, CA). For data quality control, FASTQC was used to check the raw fastq data quality and Trimmomatic was used to remove adaptors and to trim quality bases. After adapter clipping, leading and trailing ambiguous or low-quality bases were removed. The reads were then mapped to the latest UCSC transcript set using Bowtie2 version 2.1.0 and the gene expression level was estimated using RSEM v1.2.15. DESeq2 was used to find differentially expressed genes in *Tl1a-tg* compared to WT crypts. RNA-Seq data are publicly available in the NCBI Gene Expression Omnibus under GSE228942.

### Electron transmission microscopy

Ileal tissue fragments from age- and gender-matched WT and *Tl1a-tg* mice were handled by standard methods to be fixed with 2.5% glutaraldehyde, 2.0% paraformaldehyde in 0.1 M sodium cacodylate buffer, pH 7.4. After buffer washes, the samples were post-fixed with 1% OsO4 in 0.1 M sodium cacodylate for 1 h at room temperature, rinsed in dH2O and incubated with 2% uranyl acetate at RT for 1 h. After dehydration through a graded ethanol series, samples were infiltrated with Epon resin and embedded in freshly prepared Epon resin. After polymerization at 60°C for 48 h, samples were sectioned and placed on copper grids. 75-nm-thick sections were stained with uranyl acetate and lead citrate and examined with a JEOL 100CX transmission electron microscope at 60 kV operating voltage. Images were captured by an AMT digital camera (Advanced Microscopy Techniques Corporation, model XR611) with the assistance of the University of California Los Angeles Electron Microscopy Core Facility (UCLA Brain Research Institute).

### Terminal deoxynucleotidyl transferase-dUTP nick end labeling (TUNEL) assay

For cell death detection, TUNEL assay was performed using Click-iT Plus TUNEL assay kit according to manufacturer’s instruction (C10618, Thermo Fisher Scientific). Fixed ileal tissues from WT and *Tl1a-tg* mice were used.

### Gene expression analysis and quantitative trait association in human small bowel tissue sections

Transcriptomic data was generated using Whole Human Genome 4×44K Microarrays (Agilent-014850) from formalin-fixed paraffin-embedded tissue taken from the unaffected margin of small bowel tissue resected during small bowel resection for complicated CD (32). The degree of Paneth cell abnormalities was scored in lysozyme-stained slides by a trained pathologist using an established Paneth cell scoring system for human patients (D0-normal, D1-4 various degree of abnormalities) (29). Mucosal scrapings from the same slides were used to perform global gene expression analysis. The methods used to process microarray and genotyping data for the SB139 cohort have been described previously (32). Paneth cell phenotype data were available for 132 out of 139 CD patients.

### Generation and culturing of human inducible Pluripotent Stem Cell (iPSC) lines

iPSC line CS03iCTR-n1 was obtained from the iPSC Core at Cedars-Sinai and was derived from a healthy control subject. This line was fully characterized and confirmed to be karyotypically normal. It was maintained in an undifferentiated state on Matrigel-coated plates in mTeSR1 media (Stem Cell Technologies) under feeder-free conditions.

### Generation of iPSC-derived human intestinal organoids (HIOs) and Paneth cell Chip

The epithelial component of iPSC-derived HIOs was incorporated into small micro-engineered as previously described (33). Briefly, iPSCs were directed to form definitive endoderm, hindgut structures and ultimately organoids and were then overlaid with Advanced Dulbecco’s Modified Eagle Medium/F-12 with penicillin/streptomycin and L-glutamine (5% v/v) containing CHIR99021 (2 µM; Tocris), noggin, EGF (both 100 ng/ml; all R&D Systems), and B27 (1 x; Invitrogen). HIOs were then dissociated to a single cell suspension, incubated with CD326 (EpCAM) MicroBeads (Miltenyi Biotec) for 30 minutes at 4 °C, and EpCAM^+^ cells were obtained via MACS. 2 x 10^5^ EpCAM^+^ cells were subsequently incorporated into small micro-engineered Chips (Emulate) and were subjected to continuous media flow (30 μl/h) using HIO medium supplemented with SB202190 (10 µM; Tocris), and A83-01 (500 nM; Tocris) for four days. To drive Paneth cell development, medium was replaced with HIO medium containing DAPT (10 μM, Tocris) and cultured for additional four days. TL1A (10 ng/ml) was added on day four and all Chips were harvested on day eight.

### Culture of ileal organoids, and Paneth cell enrichment

Ileal organoid culture was performed using ileal crypts isolated from indicated mice as described previously (53) (54). On day 1, organoids were treated with TL1A (1896, R & D Systems). RNA from organoids was extracted by adding RLT lysis buffer (Qiagen) directly to the organoids on day 6 of culture. Paneth cell enrichment in organoids was performed as described (34, 55, 56). In brief, on day 3 of organoid culture fresh ENR media supplemented with 3 µM CHIR99021 (SML1046, Millipore Sigma) and 10 µM DAPT (D5942, Millipore Sigma) (ENR-CD) was added. On day 5, fresh ENR-CD media with or without 100 ng/ml TL1A was added. After 48 h of TL1A treatment, organoids were harvested using gentle cell dissociation reagent for immunofluorescence staining or directly lysed in RLT buffer. RNA was extracted with using RNeasy micro kit (Qiagen).

### Isolation of mesenteric lymph nodes and lamina propria mononuclear cells, restimulation, flow cytometry, and ELISA

Mesenteric lymph nodes (MLN) were removed from the peritoneal cavity and mesenteric fat was removed. Single cell suspension was prepared by crushing MLN between two glass slides and passing the cell suspension through a 40 μm cell strainer. Colons were swiftly excised and opened longitudinally followed by washing with PBS. Epithelial cells were scraped off with a glass slide and collected for RNA isolation or protein extraction. Lamina propria mononuclear cells (LPMC) were isolated as previously described (10). Single-cell suspensions of LPMC and MLN were restimulated with anti-CD3ε (0.5 μg/ml) and anti-CD28 (1 μg/ml) antibodies for 3 days. Cytokine levels in supernatants were measured by ELISA. Cytokine concentration in culture supernatants was assayed by ELISA for murine IFN-γ (eBioscience). For intracellular staining, cells were re-stimulated with 50 ng/ml phorbol 12-myristate 13-acetate (PMA), 500 ng/ml ionomycin, and Monensin (cat. # 00-4505, Thermo Fisher Scientific) for 4 h, stained with Live/dead dye (Thermo Fisher Scientific) and anti-CD4, fixed and permeabilized using the FoxP3 staining buffer set (Thermo Fisher Scientific) and stained with antibodies against murine IL-17A, IFN-γ, RORγt, Ki67, and FoxP3. For surface staining, cells were stained with Live/dead dye, and antibodies against murine CD4, CD44, and CD62L. Samples were acquired using a BD FACSymphony A5 cell analyzer (BD Biosciences, Franklin Lakes, NJ) or Sony ID7000™ Spectral Cell Analyzer and analyzed using FlowJo software (TreeStar Inc., Ashland, OR).

### Statistics

Data are presented as means ± SD. Differences between groups were calculated using two-tailed Student *t*-test for two groups and by 1-way ANOVA with Tukey’s HSD test for more than two groups. Differences were considered significant at *p* < 0.05.

### Study approvals

All mice were handled according to the guidelines and approved protocols of the institutional animal care and use committee (protocol: 8793). iPSC lines and protocols used in this study were carried out in accordance with the guidelines approved by the stem cell research oversight committee and institutional review board at Cedars-Sinai Medical Center under the auspice of the institutional review board stem cell research oversight committee protocol Pro00027264 (Derivation of Intestinal Stem Cells). All patients provided written informed consent approved by the Cedars-Sinai Medical Center institutional review board (IRB) (protocol 3358).

## Data availability

All data associated with this study are available in the main text or the supplementary materials. The datasets generated and analyzed in the current study are publicly available (NCBI Gene Expression Omnibus: GSE228942; NCBI BioProject: PRJNA951424).

## Acknowledgements

We appreciate the support received from the Cedars-Sinai Medical Center Confocal Microscopy Core, Biobank and Translational Research Core, Flow Cytometry Core, the Electron Microscopy Core Facility at the UCLA Brain Research Institute, Microbiome Core of the Goodman-Luskin Microbiome Center at UCLA, Technology Center for Genomics & Bioinformatics at UCLA, West Coast Metabolomics Center at UC Davis, and the National Gnotobiotic Rodent Resource Center at the University of North Carolina, Chapel Hill. We thank the MIRIAD IBD Biobank for providing specimen and medical data. The MIRIAD IBD Biobank is supported by the F. Widjaja IBD Institute, National Institute of Diabetes and Digestive and Kidney Disease Grant U01DK062413, and The Leona M. and Harry B. Helmsley Charitable Trust. Defa6-Cre mice were kindly provided by Richard Blumberg (Brigham and Women’s Hospital, Boston, MA).

## Funding

National Institutes of Health R01 DK056328, R01 DK123511 (SRT, DQS, KSM, RJB)

National Institutes of Health K08 Career Development Award DK093578 (DQS)

National Institutes of Health P40OD010995, P01DK094779 (RBS)

National Institutes of Health R01 DK088199 (RSB)

US Department of Veterans Affairs Career Development Award IK2CX002157 (NJ) and IK2CX001717 (JPJ)

F. Widjaja Foundation (NJ, SRT, DQS, and KSM)

Chinese Government Scholarship (YY)

Crohn’s & Colitis Foundation (RBS)

## Authorship Contributions

Conceptualization: YY, NJ, DQS, SRT, KSM

Methodology: RSB

Investigation: YY, SKM, YS, EEA, HH, AYB, DTS, JHM, JPA, LST, SLC, HQE, KK, AAP, TH, EM, TCL, KW, DPBM, RBS, DQS, RJB, NJ, JPJ, KSM

Funding acquisition: SRT, KSM

Project administration: SRT, KSM

Supervision: NJ, RJB, SRT, KSM

Writing – original draft: YY, KSM

Writing – review & editing: YY, NJ, JPJ, RBS, SRT, KSM

## Patient and public involvement

Patients and/or the public were not involved in the design, or conduct, or reporting, or dissemination plans of this research.

## References

1. Graham DB, and Xavier RJ. Pathway paradigms revealed from the genetics of inflammatory bowel disease. Nature. 2020;578(7796):527-39.

2. Cordero RY, Cordero JB, Stiemke AB, Datta LW, Buyske S, Kugathasan S, et al. Trans-ancestry, Bayesian meta-analysis discovers 20 novel risk loci for inflammatory bowel disease in an African American, East Asian and European cohort. Hum Mol Genet. 2023;32(5):873–82.

3. de Lange KM, Moutsianas L, Lee JC, Lamb CA, Luo Y, Kennedy NA, et al. Genome-wide association study implicates immune activation of multiple integrin genes in inflammatory bowel disease. Nat Genet. 2017;49(2):256–61.

4. Yamazaki K, McGovern D, Ragoussis J, Paolucci M, Butler H, Jewell D, et al. Single nucleotide polymorphisms in TNFSF15 confer susceptibility to Crohn’s disease. Human molecular genetics. 2005;14(22):3499–506.

5. Michelsen KS, Thomas LS, Taylor KD, Yu QT, Mei L, Landers CJ, et al. IBD-associated TL1A gene (TNFSF15) haplotypes determine increased expression of TL1A protein. PLoS One. 2009;4(3):e4719.

6. Barrett R, Zhang X, Koon HW, Vu M, Chang JY, Yeager N, et al. Constitutive TL1A expression under colitogenic conditions modulates the severity and location of gut mucosal inflammation and induces fibrostenosis. Am J Pathol. 2012;180(2):636–49.

7. Shih DQ, Barrett R, Zhang X, Yeager N, Koon HW, Phaosawasdi P, et al. Constitutive TL1A (TNFSF15) expression on lymphoid or myeloid cells leads to mild intestinal inflammation and fibrosis. PLoS One. 2011;6(1):e16090.

8. Meylan F, Song YJ, Fuss I, Villarreal S, Kahle E, Malm IJ, et al. The TNF-family cytokine TL1A drives IL-13-dependent small intestinal inflammation. Mucosal Immunol. 2011;4(2):172–85.

9. Sands B, Peyrin-Biroulet L, Danese S, Rubin DT, Vermeire S, Laurent O, et al. OP40 PRA023 Demonstrated Efficacy and Favorable Safety as Induction Therapy for Moderately to Severely Active UC: Phase 2 ARTEMIS-UC Study Results. Journal of Crohn’s and Colitis. 2023;17(Supplement_1):i56–i9.

10. Takedatsu H, Michelsen KS, Wei B, Landers CJ, Thomas LS, Dhall D, et al. TL1A (TNFSF15) regulates the development of chronic colitis by modulating both T-helper 1 and T-helper 17 activation. Gastroenterology. 2008;135(2):552–67.

11. Castellanos JG, Woo V, Viladomiu M, Putzel G, Lima S, Diehl GE, et al. Microbiota-Induced TNF-like Ligand 1A Drives Group 3 Innate Lymphoid Cell-Mediated Barrier Protection and Intestinal T Cell Activation during Colitis. Immunity. 2018;49(6):1077–89 e5.

12. Jacob N, Jacobs JP, Kumagai K, Ha CWY, Kanazawa Y, Lagishetty V, et al. Inflammation-independent TL1A-mediated intestinal fibrosis is dependent on the gut microbiome. Mucosal Immunol. 2018;11(5):1466–76.

13. Adolph TE, Tomczak MF, Niederreiter L, Ko HJ, Bock J, Martinez-Naves E, et al. Paneth cells as a site of origin for intestinal inflammation. Nature. 2013;503(7475):272-6.

14. Cadwell K, Liu JY, Brown SL, Miyoshi H, Loh J, Lennerz JK, et al. A key role for autophagy and the autophagy gene Atg16l1 in mouse and human intestinal Paneth cells. Nature. 2008;456(7219):259-63.

15. Kaser A, Lee AH, Franke A, Glickman JN, Zeissig S, Tilg H, et al. XBP1 links ER stress to intestinal inflammation and confers genetic risk for human inflammatory bowel disease. Cell. 2008;134(5):743–56.

16. Bevins CL, Stange EF, and Wehkamp J. Decreased Paneth cell defensin expression in ileal Crohn’s disease is independent of inflammation, but linked to the NOD2 1007fs genotype. Gut. 2009;58(6):882–3; discussion 3-4.

17. Liu TC, Gurram B, Baldridge MT, Head R, Lam V, Luo C, et al. Paneth cell defects in Crohn’s disease patients promote dysbiosis. JCI Insight. 2016;1(8):e86907.

18. Lloyd-Price J, Arze C, Ananthakrishnan AN, Schirmer M, Avila-Pacheco J, Poon TW, et al. Multi-omics of the gut microbial ecosystem in inflammatory bowel diseases. Nature. 2019;569(7758):655-62.

19. Liu TC, Kern JT, Jain U, Sonnek NM, Xiong S, Simpson KF, et al. Western diet induces Paneth cell defects through microbiome alterations and farnesoid X receptor and type I interferon activation. Cell host & microbe. 2021;29(6):988–1001 e6.

20. Franzosa EA, Sirota-Madi A, Avila-Pacheco J, Fornelos N, Haiser HJ, Reinker S, et al. Gut microbiome structure and metabolic activity in inflammatory bowel disease. Nat Microbiol. 2019;4(2):293–305.

21. Zhou H, Zhou SY, Gillilland M, 3rd, Li JY, Lee A, Gao J, et al. Bile acid toxicity in Paneth cells contributes to gut dysbiosis induced by high-fat feeding. JCI Insight. 2020;5(20).

22. Cadwell K, Patel KK, Maloney NS, Liu TC, Ng AC, Storer CE, et al. Virus-plus-susceptibility gene interaction determines Crohn’s disease gene Atg16L1 phenotypes in intestine. Cell. 2010;141(7):1135–45.

23. Liu TC, Kern JT, VanDussen KL, Xiong S, Kaiko GE, Wilen CB, et al. Interaction between smoking and ATG16L1T300A triggers Paneth cell defects in Crohn’s disease. J Clin Invest. 2018;128(11):5110–22.

24. Bel S, Pendse M, Wang Y, Li Y, Ruhn KA, Hassell B, et al. Paneth cells secrete lysozyme via secretory autophagy during bacterial infection of the intestine. Science. 2017;357(6355):1047-52.

25. Chiang HY, Lu HH, Sudhakar JN, Chen YW, Shih NS, Weng YT, et al. IL-22 initiates an IL-18-dependent epithelial response circuit to enforce intestinal host defence. Nat Commun. 2022;13(1):874.

26. Lin X, Gaudino SJ, Jang KK, Bahadur T, Singh A, Banerjee A, et al. IL-17RA-signaling in Lgr5(+) intestinal stem cells induces expression of transcription factor ATOH1 to promote secretory cell lineage commitment. Immunity. 2022;55(2):237–53 e8.

27. Gaudino SJ, Beaupre M, Lin X, Joshi P, Rathi S, McLaughlin PA, et al. IL-22 receptor signaling in Paneth cells is critical for their maturation, microbiota colonization, Th17-related immune responses, and anti-Salmonella immunity. Mucosal immunology. 2020.

28. Farin HF, Karthaus WR, Kujala P, Rakhshandehroo M, Schwank G, Vries RG, et al. Paneth cell extrusion and release of antimicrobial products is directly controlled by immune cell-derived IFN-gamma. The Journal of experimental medicine. 2014;211(7):1393–405.

29. VanDussen KL, Liu TC, Li D, Towfic F, Modiano N, Winter R, et al. Genetic variants synthesize to produce paneth cell phenotypes that define subtypes of Crohn’s disease. Gastroenterology. 2014;146(1):200–9.

30. Stahl M, Tremblay S, Montero M, Vogl W, Xia L, Jacobson K, et al. The Muc2 mucin coats murine Paneth cell granules and facilitates their content release and dispersion. American journal of physiology Gastrointestinal and liver physiology. 2018;315(2):G195–G205.

31. Liu B, Gulati AS, Cantillana V, Henry SC, Schmidt EA, Daniell X, et al. Irgm1-deficient mice exhibit Paneth cell abnormalities and increased susceptibility to acute intestinal inflammation. Am J Physiol Gastrointest Liver Physiol. 2013;305(8):G573–84.

32. Potdar AA, Li D, Haritunians T, VanDussen KL, Fiorino MF, Liu TC, et al. Ileal Gene Expression Data from Crohn’s Disease Small Bowel Resections Indicate Distinct Clinical Subgroups. J Crohns Colitis. 2019;13(8):1055–66.

33. Workman MJ, Gleeson JP, Troisi EJ, Estrada HQ, Kerns SJ, Hinojosa CD, et al. Enhanced Utilization of Induced Pluripotent Stem Cell-Derived Human Intestinal Organoids Using Microengineered Chips. Cell Mol Gastroenterol Hepatol. 2018;5(4):669–77 e2.

34. Yin X, Farin HF, van Es JH, Clevers H, Langer R, and Karp JM. Niche-independent high-purity cultures of Lgr5+ intestinal stem cells and their progeny. Nat Methods. 2014;11(1):106–12.

35. Atarashi K, Tanoue T, Oshima K, Suda W, Nagano Y, Nishikawa H, et al. Treg induction by a rationally selected mixture of Clostridia strains from the human microbiota. Nature. 2013;500(7461):232-6.

36. Thorburn AN, McKenzie CI, Shen S, Stanley D, Macia L, Mason LJ, et al. Evidence that asthma is a developmental origin disease influenced by maternal diet and bacterial metabolites. Nat Commun. 2015;6:7320.

37. Gunther C, Ruder B, Stolzer I, Dorner H, He GW, Chiriac MT, et al. Interferon Lambda Promotes Paneth Cell Death Via STAT1 Signaling in Mice and Is Increased in Inflamed Ileal Tissues of Patients With Crohn’s Disease. Gastroenterology. 2019;157(5):1310–22 e13.

38. 38. Steenwinckel V, Louahed J, Lemaire MM, Sommereyns C, Warnier G, McKenzie A, et al. IL-9 promotes IL-13-dependent paneth cell hyperplasia and up-regulation of innate immunity mediators in intestinal mucosa. Journal of immunology (Baltimore, Md : 1950). 2009;182(8):4737–43.

39. Tsuda M, Hamade H, Thomas LS, Salumbides BC, Potdar AA, Wong MH, et al. A role for BATF3 in TH9 differentiation and T-cell-driven mucosal pathologies. Mucosal Immunology. 2019.

40. Imhann F, Vich Vila A, Bonder MJ, Fu J, Gevers D, Visschedijk MC, et al. Interplay of host genetics and gut microbiota underlying the onset and clinical presentation of inflammatory bowel disease. Gut. 2018;67(1):108–19.

41. Gevers D, Kugathasan S, Denson LA, Vazquez-Baeza Y, Van Treuren W, Ren B, et al. The treatment-naive microbiome in new-onset Crohn’s disease. Cell host & microbe. 2014;15(3):382–92.

42. Riba A, Olier M, Lacroix-Lamande S, Lencina C, Bacquie V, Harkat C, et al. Paneth Cell Defects Induce Microbiota Dysbiosis in Mice and Promote Visceral Hypersensitivity. Gastroenterology. 2017;153(6):1594–606 e2.

43. Wahida A, Muller M, Hiergeist A, Popper B, Steiger K, Branca C, et al. XIAP restrains TNF-driven intestinal inflammation and dysbiosis by promoting innate immune responses of Paneth and dendritic cells. Sci Immunol. 2021;6(65):eabf7235.

44. Smith PM, Howitt MR, Panikov N, Michaud M, Gallini CA, Bohlooly YM, et al. The microbial metabolites, short-chain fatty acids, regulate colonic Treg cell homeostasis. *Science (New York*, NY). 2013;341(6145):569-73.

45. Park J, Kim M, Kang SG, Jannasch AH, Cooper B, Patterson J, et al. Short-chain fatty acids induce both effector and regulatory T cells by suppression of histone deacetylases and regulation of the mTOR-S6K pathway. Mucosal immunology. 2015;8(1):80–93.

46. Raetz M, Hwang SH, Wilhelm CL, Kirkland D, Benson A, Sturge CR, et al. Parasite-induced TH1 cells and intestinal dysbiosis cooperate in IFN-gamma-dependent elimination of Paneth cells. Nature immunology. 2013;14(2):136–42.

47. Abbas AK, Trotta E, D RS, Marson A, and Bluestone JA. Revisiting IL-2: Biology and therapeutic prospects. Sci Immunol. 2018;3(25).

48. Jacob N, Jacobs JP, Kumagai K, Ha CWY, Kanazawa Y, Lagishetty V, et al. Inflammation-independent TL1A-mediated intestinal fibrosis is dependent on the gut microbiome. Mucosal Immunol. 2018.

49. Ostanin DV, Pavlick KP, Bharwani S, D’Souza D, Furr KL, Brown CM, et al. T cell-induced inflammation of the small and large intestine in immunodeficient mice. Am J Physiol Gastrointest Liver Physiol. 2006;290(1):G109–19.

50. Erben U, Loddenkemper C, Doerfel K, Spieckermann S, Haller D, Heimesaat MM, et al. A guide to histomorphological evaluation of intestinal inflammation in mouse models. Int J Clin Exp Pathol. 2014;7(8):4557–76.

51. Jacobs JP, Lin L, Goudarzi M, Ruegger P, McGovern DP, Fornace AJ, Jr., et al. Microbial, metabolomic, and immunologic dynamics in a relapsing genetic mouse model of colitis induced by T-synthase deficiency. Gut Microbes. 2017;8(1):1–16.

52. Dekkers JF, Alieva M, Wellens LM, Ariese HCR, Jamieson PR, Vonk AM, et al. High-resolution 3D imaging of fixed and cleared organoids. Nat Protoc. 2019;14(6):1756–71.

53. Sato T, Vries RG, Snippert HJ, van de Wetering M, Barker N, Stange DE, et al. Single Lgr5 stem cells build crypt-villus structures in vitro without a mesenchymal niche. Nature. 2009;459(7244):262-5.

54. Shimodaira Y, More SK, Hamade H, Blackwood AY, Abraham JP, Thomas LS, et al. DR3 Regulates Intestinal Epithelial Homeostasis and Regeneration After Intestinal Barrier Injury. Cell Mol Gastroenterol Hepatol. 2023;16(1):83–105.

55. Mead BE, Ordovas-Montanes J, Braun AP, Levy LE, Bhargava P, Szucs MJ, et al. Harnessing single-cell genomics to improve the physiological fidelity of organoid-derived cell types. BMC Biol. 2018;16(1):62.

56. Treveil A, Sudhakar P, Matthews ZJ, Wrzesinski T, Jones EJ, Brooks J, et al. Regulatory network analysis of Paneth cell and goblet cell enriched gut organoids using transcriptomics approaches. Mol Omics. 2020;16(1):39–58.

