## Supplemental data for "TL1A overexpression in Crohn’s Disease and mice alters Paneth cells and microbiota promoting ileal inflammation"

### Supplemental Figure 1

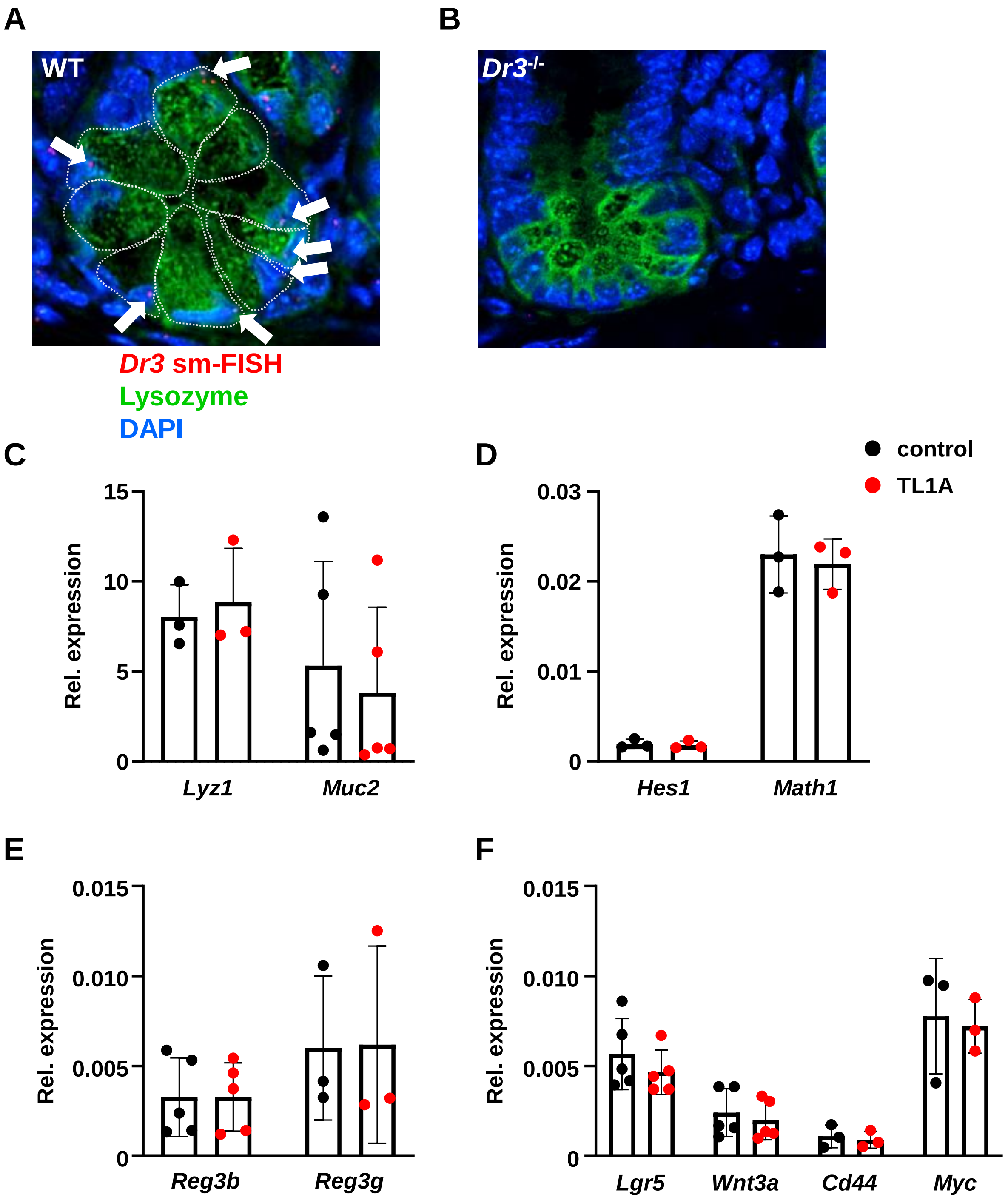


**Supplemental Figure 1.** *Dr3 is expressed on murine ileal Paneth cells.* (**A**) *Dr3* expression on ileal Paneth cells was detected by single-molecule fluorescent *in situ* hybridization (smFISH). Ileal tissue sections of WT small intestine were hybridized with smFISH probes against *Dr3* (red), stained with anti-lysozyme antibody (green), and counterstained with DAPI (blue). Expression of *Dr3* by Paneth cells is indicated by arrows. (**B**) *Dr3*$^{-/-}$ mice do not express *Dr3*. (**C - F**) Mouse ileal organoids were cultured under Paneth cell-enrichment conditions with or without TL1A. mRNA expression of *Lyz1, Muc2* (**C**), *Hes1, Math1* (**D**), *Reg3b, Reg3g* (**E**), *Lgr5, Wnt3a, Cd44, Myc* (**F**). Data represent means ± SD.

### Supplemental Figure 2

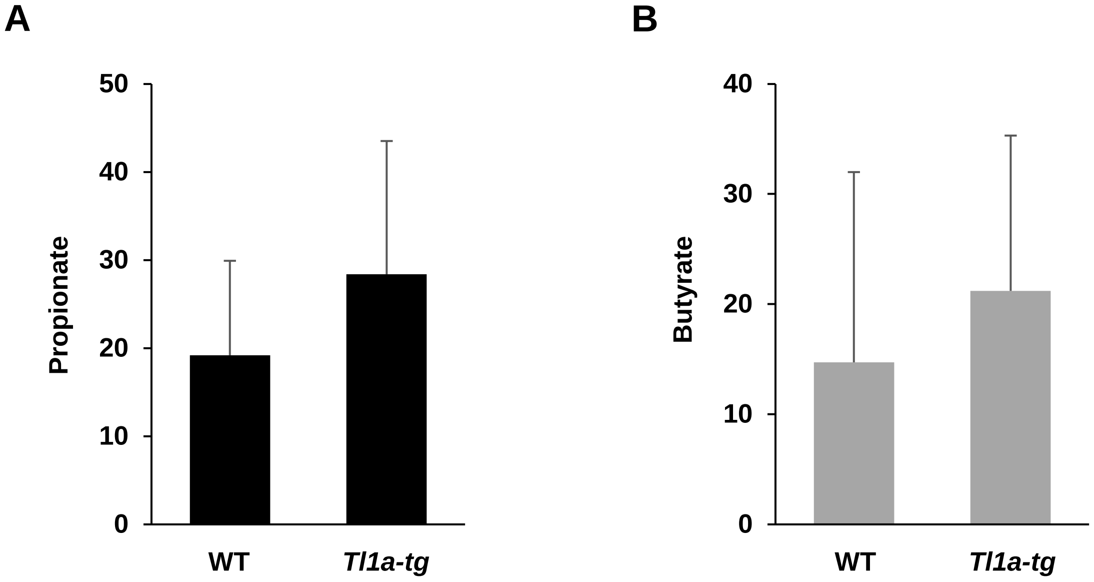


**Supplemental Figure 2. Short chain fatty acid profiling of mucosal ileal washings.** (**A - B**) Metabolomic profiling was performed of mucosal ileal washings from 2 months old SPF littermate controls (n = 10/group). (**A**) Abundance of propionate in the ileum of SPF WT and *Tl1a-tg* mice. (B) Abundance of butyrate in the ileum of SPF WT and *Tl1a-tg* mice.

### Supplemental Figure 3

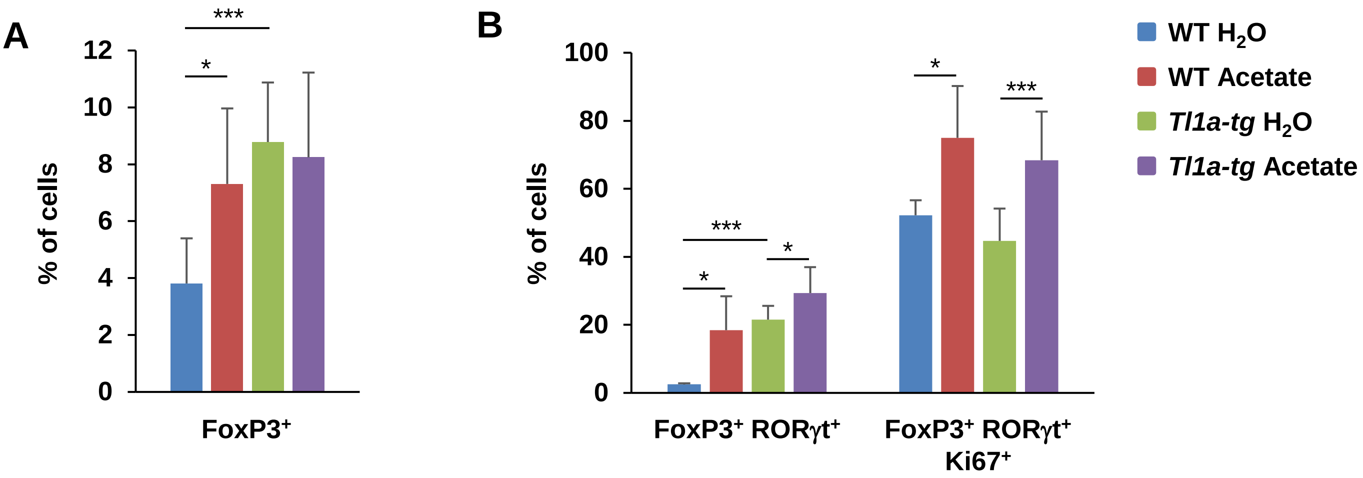


**Supplemental Figure 3. Acetate supplementation of GF mice leads to expansion of Tregs.** (**A, B**) GF mice were supplemented with acetate drinking water for 3 months. (**A, B**) Flow cytometry analysis of $CD4^+$ Treg populations in MLN (n=4-7/group). (**A**) Percent of $CD4^+$ $FoxP3^+$ cells. (**B**) Percent of $CD4^+$ $FoxP3^+$ $ROR\gamma t^+$ and $CD4^+$ $FoxP3^+$ $ROR\gamma t^+$ $Ki67^+$ cells. *$p<0.05$, *** $p<0.005$ by Student's *t*-test.

### Supplemental Figure 4

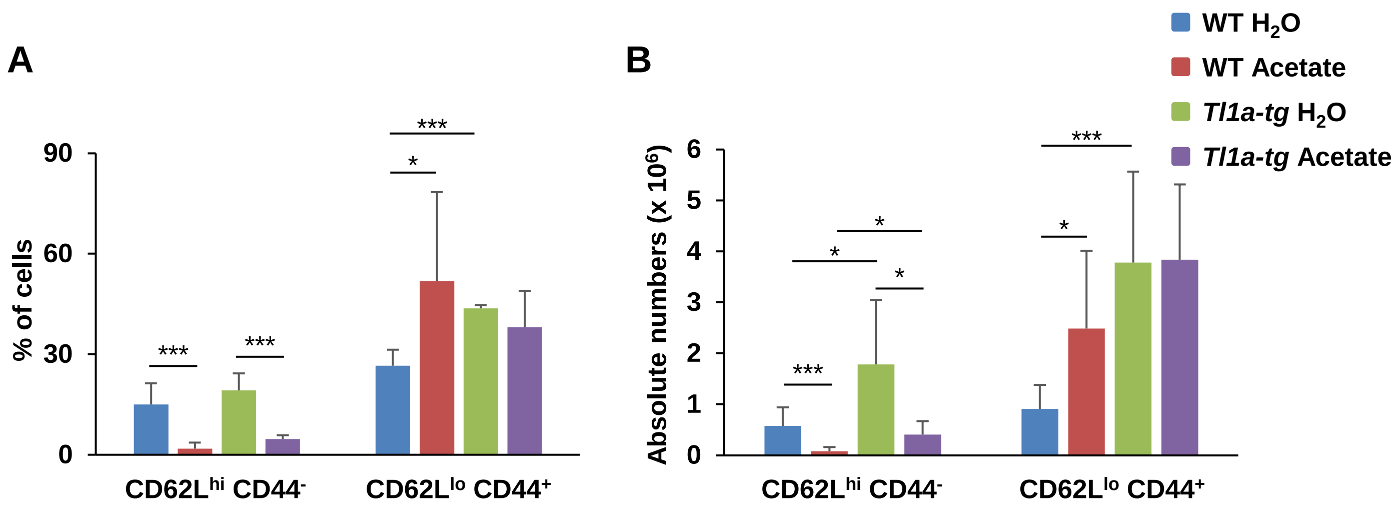


**Supplemental Figure 4. Acetate supplementation of SPF mice leads to an increase of effector T cells.** (**A, B**) SPF mice were supplemented with acetate drinking water for 2 months. (**A, B**) Flow cytometry analysis of MLN cells (n = 2-8/group). (**A**) Percent of CD4$^{+}$ naïve T cells (CD62L$^{hi}$ CD44$^{-}$) and CD4$^{+}$ effector T cells (CD62L$^{lo}$ CD44$^{+}$) cells. (**B**) Total numbers of CD4$^{+}$ naïve T cells (CD62L$^{hi}$ CD44$^{-}$) and CD4$^{+}$ effector T cells (CD62L$^{lo}$ CD44$^{+}$) cells. *$p<0.05$, *** $p<0.005$ by Student's *t*-test.

### Supplemental Figure 5

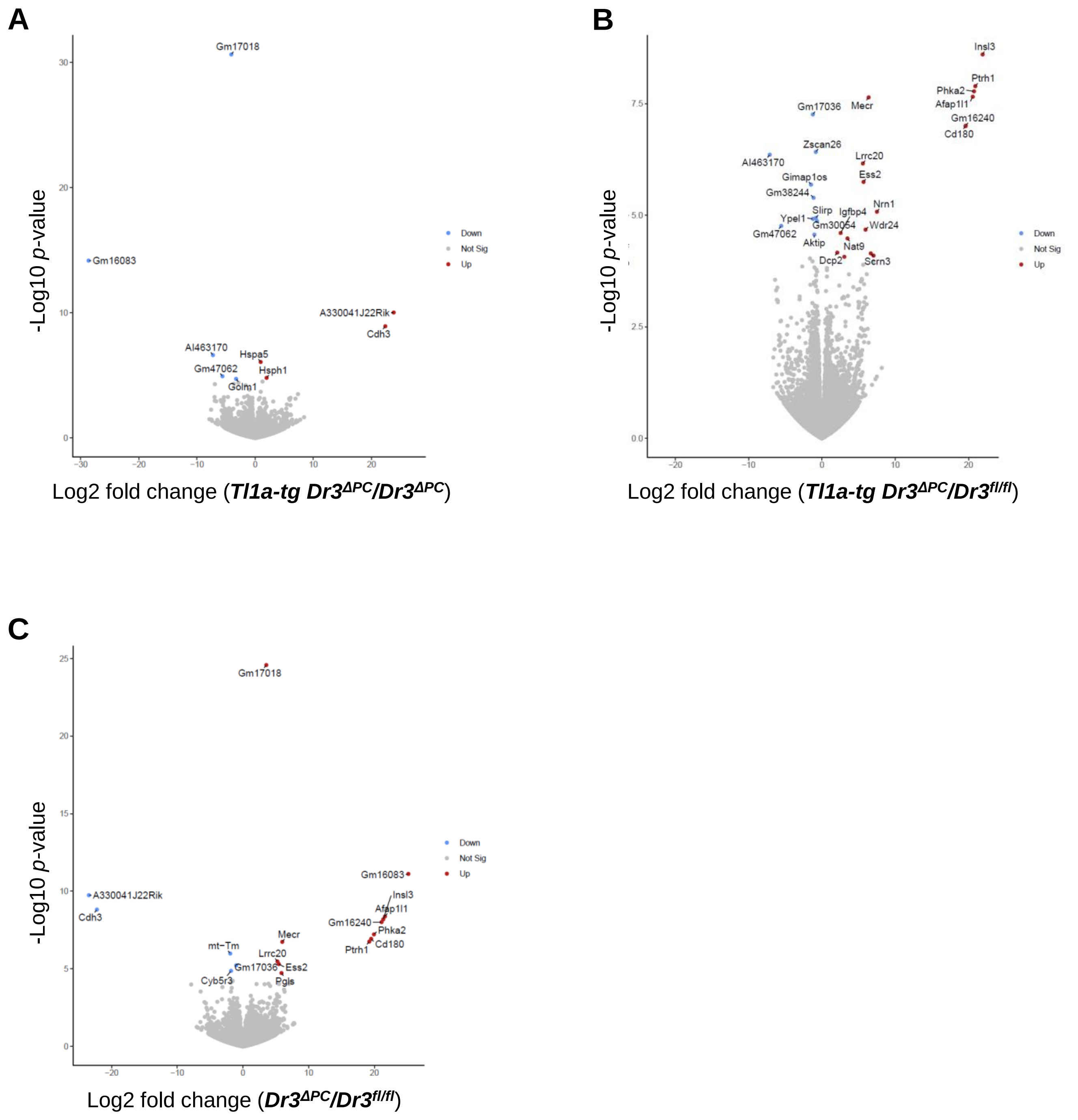


**Supplemental Figure 5. Transcriptional profiling of isolated** ***$Dr3^{fl/fl}$*****,** ***$Dr3^{\Delta PC}$*****, and** ***Tl1a-tg $Dr3^{\Delta PC}$*** **ileal crypts.** (**A-C**) Transcriptional profiling of small intestinal crypts of untreated 3-month-old *$Dr3^{fl/fl}$*, *$Dr3^{\Delta PC}$*, and *Tl1a-tg $Dr3^{\Delta PC}$* mice (n = 4 mice/group). Volcano Plots of top differentially expressed genes (DEG) with $p_{adj} < 0.05$ in small intestinal crypts from *Tl1a-tg $Dr3^{\Delta PC}$* vs. *$Dr3^{\Delta PC}$* (**A**), *Tl1a-tg $Dr3^{\Delta PC}$* vs. *$Dr3^{fl/fl}$* (**B**), or *$Dr3^{\Delta PC}$ vs. $Dr3^{fl/fl}$* (**C**) mice.

### Supplemental Figure 6

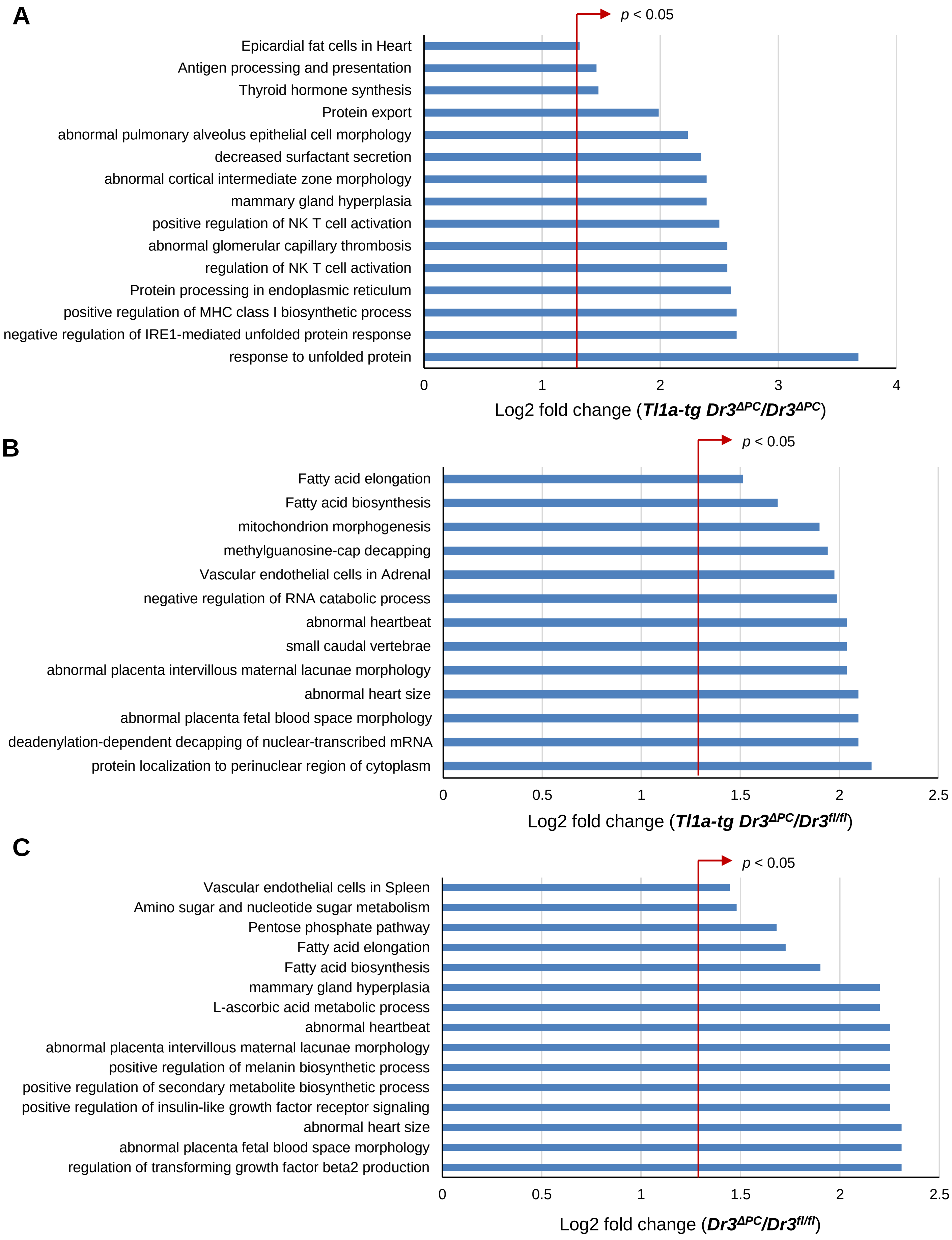

**Supplemental Figure 6. Transcriptional profiling of isolated *Dr3$^{fl/fl}$*, *Dr3$^{\Delta PC}$*, and *Tl1a-tg Dr3$^{\Delta PC}$* ileal crypts.** (**A**) Pathway and Gene Ontology analysis of DEG in *Tl1a-tg Dr3$^{\Delta PC}$* vs. *Dr3$^{\Delta PC}$* isolated small intestinal crypts. (**B**) Pathway and Gene Ontology analysis of DEG in *Tl1a-tg Dr3$^{\Delta PC}$* vs. *Dr3$^{fl/fl}$* isolated small intestinal crypts. (**C**) Pathway and Gene Ontology analysis of DEG in *Dr3$^{\Delta PC}$ vs. Dr3$^{fl/fl}$* isolated small intestinal crypts.

Supplemental Figure 7

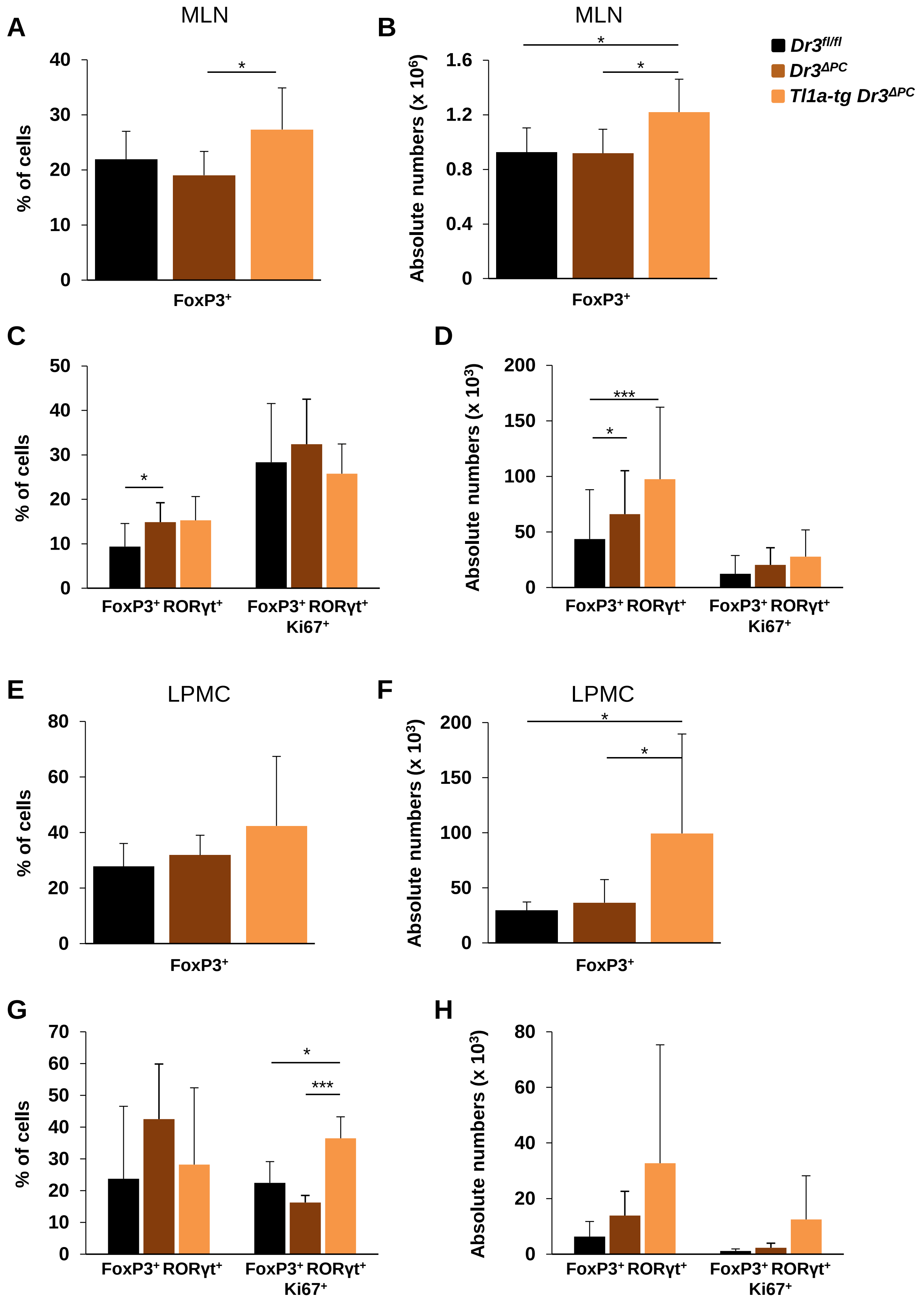

**Supplemental Figure 7. Steady-state immunoprofiling of regulatory T cells (Tregs) in *Dr3^fl/fl^*, *Dr3^ΔPC^*, and *Tl1a-tg Dr3^ΔPC^* mice.** Homeostatic MLN and lamina propria immune profiles of 6 to 12-months-old *$Dr3^{fl/fl}$*, *$Dr3^{\Delta PC}$*, and *Tl1a-tg $Dr3^{\Delta PC}$* mice were analyzed. (**A - D**) Cells were isolated from MLN and percentages (**A, C**) and absolute numbers (**B, D**) of $CD4^+$ $FoxP3^+$ , $CD4^+$ $FoxP3^+$ $ROR\gamma t^+$, $CD4^+$ $FoxP3^+$ $ROR\gamma t^+$ $Ki67^+$ cells are shown (n = 6 - 8/genotype). (**E - H**) Cells were isolated from large intestinal lamina propria and percentages (**E, G**) and absolute numbers (**F, H**) of $CD4^+$ $FoxP3^+$ , $CD4^+$ $FoxP3^+$ $ROR\gamma t^+$, $CD4^+$ $FoxP3^+$ $ROR\gamma t^+$ $Ki67^+$ cells are shown (n = 4 - 6/genotype). Means $\pm$ SD are shown. *$p < 0.05$, ***$p < 0.005$; Student's *t*-test.

### Supplemental Figure 8

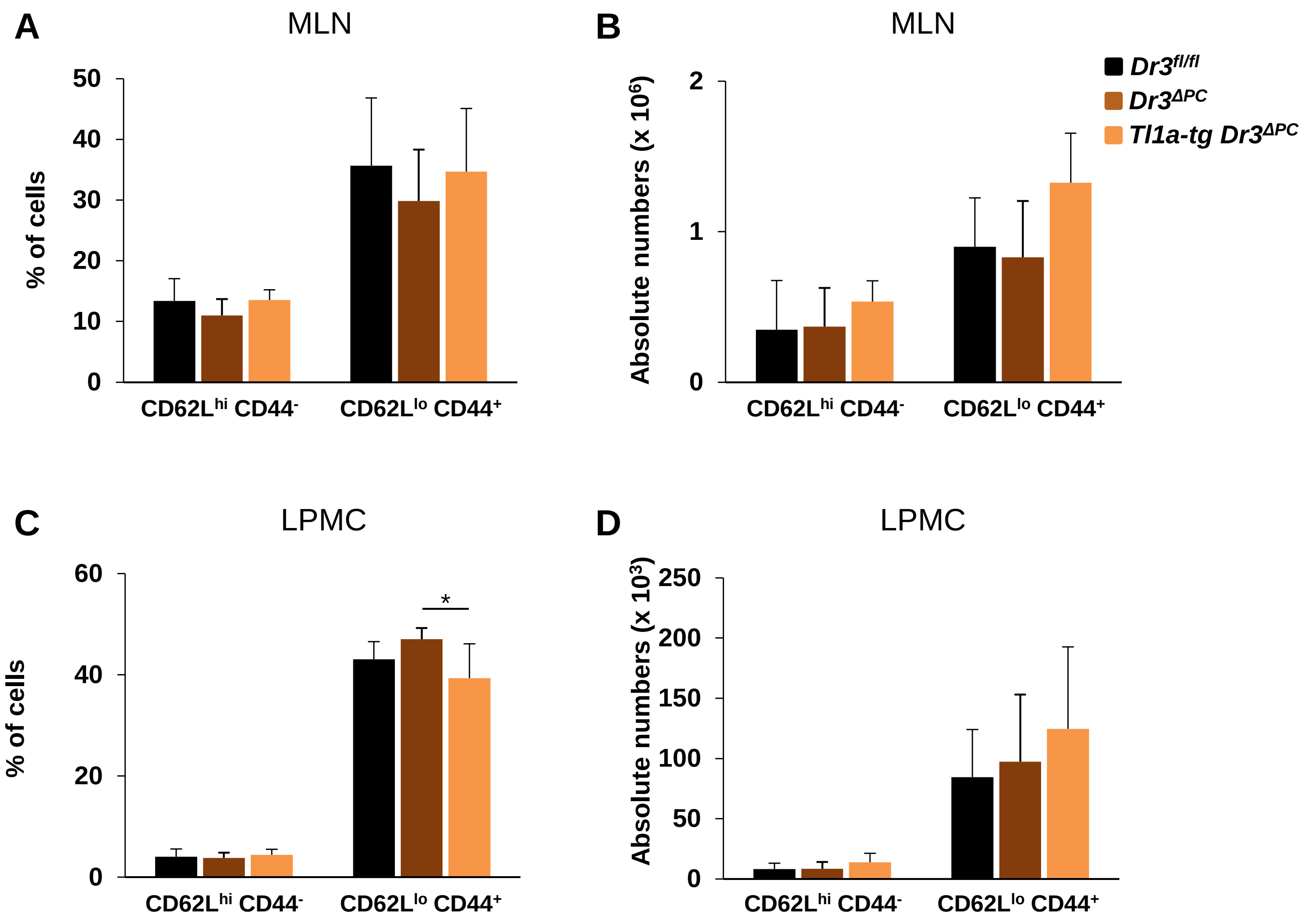


**Supplemental Figure 8. Steady-state immunoprofiling of naïve and effector CD4$^{+}$ T cells in *Dr3$^{fl/fl}$*, *Dr3$^{\Delta PC}$*, and *Tl1a-tg Dr3$^{\Delta PC}$* mice.** Homeostatic MLN and lamina propria immune profiles of 6 to 12-months-old *Dr3$^{fl/fl}$*, *Dr3$^{\Delta PC}$*, and *Tl1a-tg Dr3$^{\Delta PC}$* mice were analyzed. (**A, B**) Cells were isolated from MLN and percentages (**A**) and absolute numbers (**B**) of CD4$^{+}$ CD62L$^{hi}$ CD44$^{-}$, CD4$^{+}$ CD62L$^{lo}$ CD44$^{+}$ cells are shown (n = 6 - 9/genotype). (**C, D**) Cells were isolated from large intestinal lamina propria and percentages (**C**) and absolute numbers (**D**) of CD4$^{+}$ CD62L$^{hi}$ CD44$^{-}$, CD4$^{+}$ CD62L$^{lo}$ CD44$^{+}$ cells are shown (n = 4 - 6/genotype). Means ± SD are shown. *$p < 0.05$; Student's *t*-test.
